# Mutation-selection-drift balance models of complex diseases

**DOI:** 10.1101/2025.05.18.654722

**Authors:** Jeremy J. Berg, Xinyi Li, Kellen Riall, Laura K. Hayward, Guy Sella

## Abstract

Genetic variation that influences complex disease susceptibility is introduced into the population by mutation and removed by natural selection and genetic drift. This mutation-selection-drift-balance (MSDB) shapes the prevalence of a disease and its genetic architecture. To date, however, MSDB has only been modeled for monogenic (Mendelian) diseases. Here, we develop a MSDB model for complex disease susceptibility: we assume that genotype relates to disease risk according to the canonical liability threshold model and that selection on variants affecting risk derives from the fitness cost of the disease, and focus on diseases that are highly polygenic, entail a substantial fitness cost, and are neither extremely common in the population nor exceedingly rare. Contrasting model predictions with GWAS and other findings in humans suggests that directional selection plays little role in shaping common genetic variation affecting complex disease susceptibility but might substantially affect rare, large effect variation. In turn, common variation affecting complex disease susceptibility appears to be dominated by pleiotropic stabilizing selection on other traits. Our results further suggest that current estimates of disease heritability are likely biased. More generally, our model provides a better understanding of the evolutionary processes that shape the architecture and prevalence of complex diseases.

## Introduction

A central goal of population genetics is to understand how evolutionary processes shape the prevalence of genetic diseases and the population distribution of their underlying genetic variants. This question is of particular interest in humans. Since the late 20^th^ century, we have learned a lot about the genetic basis of simple (Mendelian) diseases and the frequencies of their underlying variants in human populations (Jobling et al. 2013, Ch. 16). We also have long-standing models for the evolutionary processes that generate and maintain these diseases (Haldane 1927; Fuller et al. 2019), whose predictions are in qualitative, if not quantitative, agreement with empirical observations (see, e.g., Amorim et al. 2017).

Most common genetic diseases in humans (e.g., with prevalence ≥ 0.1%) are complex (Jobling et al. 2013. Ch. 17), however, and it is only over the past decade or so that genome-wide association studies (GWAS) have begun to reveal their genetic basis (The Wellcome Trust Case Control Consortium 2007; Trubetskoy et al. 2022). These studies have now identified many thousands of robust associations between genetic variants and many diseases, and in so doing, have begun to uncover the numbers, effect sizes and frequencies of the variants underlying disease risk—henceforth the “genetic architecture” of complex diseases (Abdellaoui et al. 2023). Yet we still lack a good understanding of the evolutionary processes that shape the architecture and prevalence of complex diseases.

The discoveries from GWAS shed some light on these processes. Notably, they reveal that variant effects on disease risk are negatively correlated with their minor allele frequency, indicating that natural selection acts to remove genetic variation affecting disease risk and that the strength of selection on variants increases with their effect on risk (Schoech et al. 2019; Zeng et al. 2021). Additionally, many of the significant associations in GWAS of diseases are common (see, e.g., Trubetskoy et al. 2022) indicating that for much of the variation affecting disease risk, the effects of selection on variant frequencies are comparable to those of random genetic drift. It further appears that variation in the risk of developing common complex diseases is thinly spread among many thousands of segregating variants that are widely distributed across the genome (Loh et al. 2015; Shi, Kichaev, and Pasaniuc 2016; Boyle, Li, and Pritchard 2017). This extreme polygenicity, alongside evidence for selection and drift that lead to the removal of variation, implies that genetic variation is continually replenished by mutations at numerous sites across the genome. Thus, complex disease prevalence and architecture are shaped by a balance between mutation, natural selection, and genetic drift.

Existing models for mutation-selection-drift balance (MSDB) come in three flavors. The first is the classic model for simple Mendelian diseases introduced by Danforth and Haldane and extended by Muller, Kimura and others (Danforth 1923; Haldane 1927; 1937; Muller 1950; Crow 1958; Kimura 1961; Kimura, Maruyama, and Crow 1963; Clark 1998). In its most basic form, the model assumes that mutations arising at a single gene cause the disease in either heterozygotes (the dominant case) or homozygotes (the recessive case) and that the disease markedly reduces individual fitness (see, e.g., Gillespie 2004). The fitness cost of the disease induces strong selection against the alleles that cause it, leading to their loss from the population. The model describes the prevalence of the disease, the frequency of the underlying alleles, and the genetic load as a function of the mutation rate and fitness cost. This model and its generalizations, however, are not applicable to complex diseases, because the risk of developing complex diseases arises from the contribution of many variants (and effects of the environment), which, in turn, generates a less obvious relationship between the fitness cost of the disease and selection on its underlying variants.

The second flavor of MSDB models relate selection on complex traits to the selection acting on the many variants that affect these traits. They do not focus on disease risk, however, but on quantitative (continuous) traits, assuming that these traits are subject to stabilizing selection, i.e., that traits have an optimal value and individual fitness declines continuously with displacement from it (Robertson 1956; Lande 1975; Keightley and Hill 1988; Simons et al. 2018). Typically, these models further assume that mutations are equally likely to increase or decrease trait values, and that the effect sizes in either direction have the same distribution, an assumption known as symmetric mutation (but see Waxman and Peck 2003; Zhang and Hill 2008; Charlesworth 2013a; 2013b). MSDB models of quantitative, complex traits have been invaluable is studying the processes that maintain heritable variation in complex traits. More recently, they have been used to study the genetic architecture of complex traits and to interpret the results of human GWAS (Simons et al. 2018; O’Connor et al. 2019; Zeng et al. 2021; Simons et al. 2022; Spence et al. 2024). It is not obvious that they apply to complex diseases, however.

Indeed, some diseases, for example hypertension or obesity, are defined in terms of underlying quantitative traits that exceed a threshold value, and these underlying traits may well be subject to stabilizing selection and have symmetric mutation. Other diseases, for instance type 2 diabetes, may reflect a discrete biological dysfunction, such as a breakdown of homeostasis (Alon 2023). In such cases, we might expect the effects of the disease on fitness to be discrete, selection to be directional, in always acting to reduce disease risk, and mutations may therefore tend to increase disease risk.

The third flavor of MSDB models includes all these elements, while considering fitness rather than disease risk as the focal trait. These models were introduced to study the fitness burden of deleterious mutations—the genetic load—in natural populations (King 1966; Kimura and Maruyama 1966; Alexey S. Kondrashov 1995) and were later related to evolutionary advantages of sex (Alexey S. Kondrashov 1982; 1988). They typically assume that fitness drops sharply around some threshold number of deleterious mutations or threshold ‘liability’ (sometimes referred to as ‘fitness potential’; Milkman 1978; Kondrashov 2018), which arises from additive (weighted) contributions over all the deleterious alleles that an individual carries. With fitness always decreasing with an increasing number of deleterious alleles (or with increasing liability), selection is always directed against these alleles. Under such directional selection, loci tend to be fixed for beneficial alleles and therefore mutations at these loci tend to be deleterious. With some modifications, these models could be framed as models of complex diseases that are similar to the model we present below.

In their existing form, however, such models cannot be related to the architecture and prevalence of complex disease. Indeed, some of the models (Kimura and Maruyama 1966; Alexey S. Kondrashov 1982; 1984) assume that the variants that underlie fitness are all subject to strong selection (i.e., selection that is much stronger than genetic drift), an assumption that contradicts what we have learned from GWAS of complex diseases (see above). Other studies rely on rough approximations to describe weakly selected variation (e.g. Kondrashov 1995) or assume a mapping between liability and fitness that is inconsistent with disease models (Charlesworth 1990; 2013b). These assumptions do not allow the genetic architecture and prevalence of complex diseases to be related to their underlying evolutionary parameters.

Motivated by these considerations, we develop a MSDB model of complex disease risk and solve it for the genetic architecture of the disease and its prevalence. Similar to classic models of simple (Mendelian) diseases and analogous to models of genetic load, we assume that directional selection always acts to reduce disease risk, and that the disease state is discrete. Similar to models of complex quantitative traits and analogous to models of genetic load, we assume that disease risk arises from the joint effects of many variants and environmental effects. Unlike models of genetic load, we consider accurate approximations for the behavior of weakly selected variation affecting disease risk. By combining these features, we are able to describe the expected genetic architecture of complex diseases and their prevalence and to contrast our predictions with the findings of GWAS in humans.

## The Model

We use the canonical model for binary traits in human and quantitative genetics—the liability threshold model—to relate an individual’s genotype with their risk of developing a disease (Wright 1926; 1934; Lush, Lamoreux, and Hazel 1948; Dempster and Lerner 1950; Falconer 1965). Liability is a quantitative (continuous) trait that cannot be observed directly. When an individual’s liability, *Z*, exceeds a threshold, *T*, they will develop or have the disease (Figure 1A).

**Figure 1.**
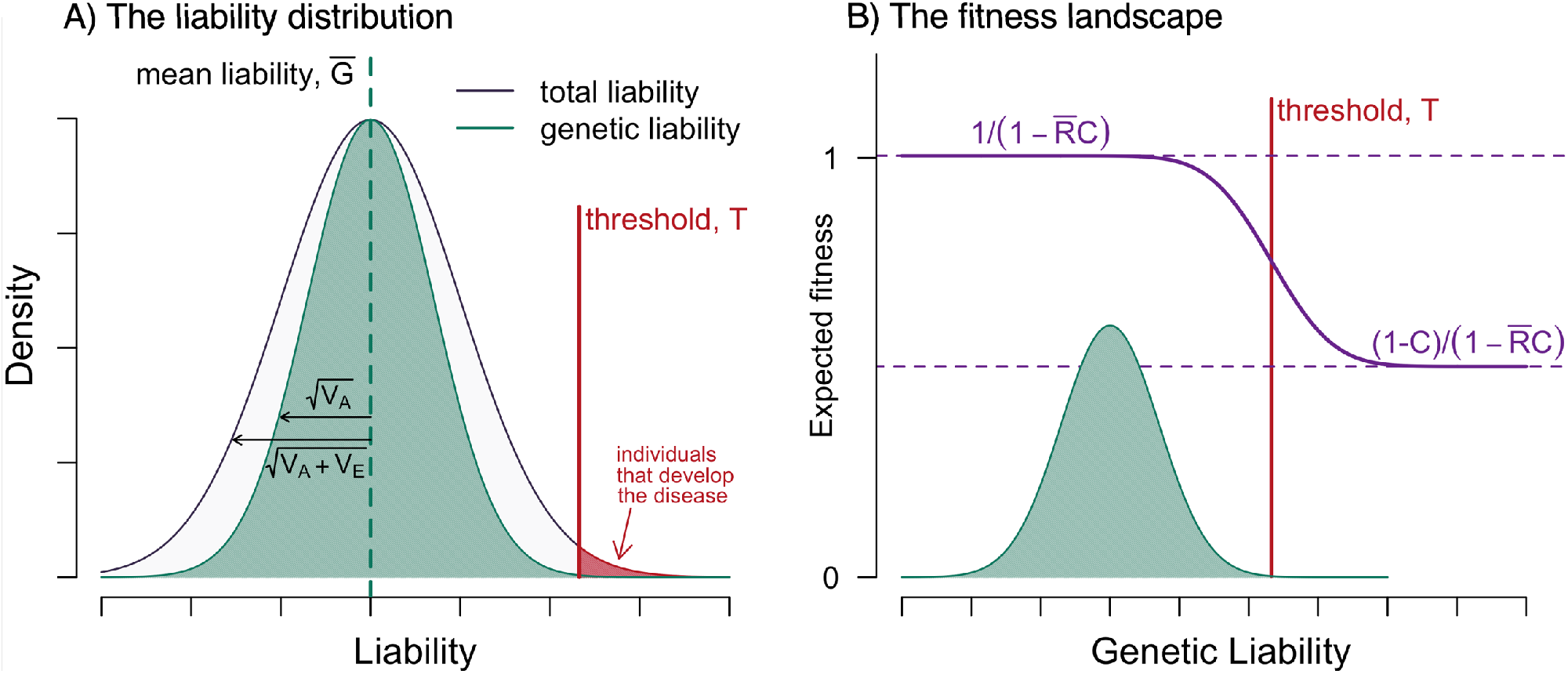
The model. (A) The liability distribution in the population. Individuals whose liability exceed the threshold develop the disease (in red). (B) The fitness landscape. In these illustrations, we assume that the genetic liability distribution is Normal, that the heritable variance in liability *h*^2^ = 1/2, the fitness cost *C* = 1/2, and the prevalence 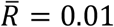.

An individual’s liability relates to their genotype through the standard additive model of quantitative complex traits (Falconer and Mackay 1995). Namely, we assume that the number of genomic sites affecting liability (i.e., the target size) is very large, *L* ≫ 1, and that an individual’s liability is given by

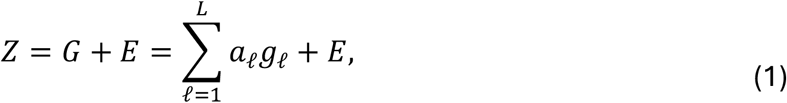

where *G* is the genetic contribution, which is the sum of contributions over sites, *a*_*𝓁*_ is the effect size of the liability increasing allele at site *𝓁*, and *g*_*𝓁*_ is the number of copies of that allele at that site; and *E*∼*N*(0, *V*_*E*_) is the environmental contribution.

This model implies that an individual’s total liability is normally distributed around their genetic liability, with the variance arising from the environmental contribution. Namely, that *Z*|*G*∼*N*(*G, V*_*E*_). An individual’s probability of developing the disease, their genetic risk, is therefore

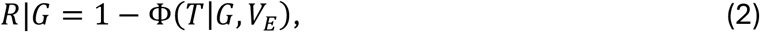

where Φ(· |*G, V*_*E*_) is the Normal cumulative distribution function with mean *G* and variance *V*_*E*_.

We model fitness by assuming that the disease entails a fitness cost *C*, such that an individual without the disease has fitness 1 and one with the disease has fitness 1 − *C*. The mean fitness of a population with disease prevalence 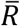 is therefore 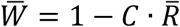, and the relative fitness associated with genetic liability *G* is

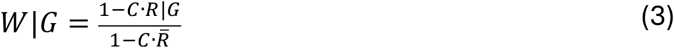

(with *R*|*G* given in Eq. 2).

Figure 1B shows the resulting fitness landscape. Fitness plateaus at 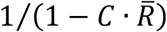 at low genetic liability and at 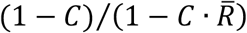 at high genetic liability, and the width of the transition between plateaus reflects the variance of the environmental contribution. This drop-off in fitness leads to selection to reduce the mean genetic liability of the population.

We assume, for simplicity, that the genomic sites affecting liability are bi-allelic. We set the liability scale such that at a given site *𝓁*, the low liability allele contributes 0 and the high liability allele contributes liability *a*_*𝓁*_ > 0. We denote the distribution of effects across sites by *g*(*a*) and its mean by 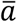. In these terms, the possible values of genetic liability range between 0 and 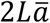 (when all sites are fixed for the low or high liability alleles, respectively).

Each generation, mutation introduces new variants that affect liability into the population. We assume that the two possible alleles mutate to one another with probability *u* per gamete per generation. We further assume, for simplicity, that the rate of mutational input per site in the population is sufficiently low for us to rely on the infinite sites approximation (specifically, we assume that *θ*/2 = 2*Nu* ≪ 1, where *N* is the population size). In this approximation, segregating derived alleles are assumed to have arisen from a single mutation, and the number of mutations per gamete per generation follows a Poisson distribution with mean *Lu*. The infinite sites approximation is standard and sensible in many contexts in humans (Harpak, Bhaskar, and Pritchard 2016; Schraiber, Spence, and Edge 2024).

The population dynamics follow the standard model of a diploid, panmictic population of constant size *N*, with non-overlapping generations. In each generation, parents are randomly chosen to reproduce with probabilities proportional to their fitness (Eq. 3), i.e., Wright-Fisher sampling with fertility selection, followed by mutation, free recombination (i.e., no linkage) and Mendelian segregation. Our notation is summarized in ***Table S1***.

### Scope

We focus on diseases that are highly polygenic, have a substantial fitness cost, and are not extremely common or exceedingly rare. Specifically, our analysis should apply to diseases with a target size *L* ≳ 10^5^ (noting that the target size is typically much greater than the polygenicity), fitness cost *C* ≳ 0.1, and prevalence 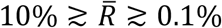. These assumptions plausibly encompass most common complex diseases in humans. Moreover, within these parameter ranges, the liability threshold model on which we rely and other standard models of complex diseases in human statistical genetics are practically interchangeable (Slatkin 2008; Wray and Goddard 2010); these include Risch’s multiplicative model used as premise in linkage studies (Risch 1990) and the logistic model used in case-control GWAS (Sham and Purcell 2014).

We focus on the model’s behavior at MSDB (i.e., at equilibrium). For the type of diseases consider, most of the liability distribution falls below the threshold, with only a small tail above it (Fig. 1A). For simplicity, we assume that the liability distribution is well approximated by its stationary distribution and ignore stochastic fluctuations around this stationary distribution. Under our assumptions (notably of high polygenicity), this is a sensible assumption that should not have a substantial effect our results.

### Simulations

We validate our analytic results using simulations. The simulations are implemented in SLiM (version 3.6; Haller and Messer 2019) and realize the models with one or two liability effect sizes. We initialize the simulations with genetically identical, homozygous individuals at all sites. We set effect sizes at sites according to their expected fixed state at MSDB, which we derive below (otherwise, reaching MSDB would take too long to be computationally feasible). We run the simulation for a burn-in period of 10*N* generations to allow genetic variation to be near the steady state at MSDB. We then run the simulation for an additional 25*N* generations and sample the population every 0.05*N* generations to collect 500 samples per simulation. We run 6 replicate simulations for each parameter setting and estimate quantities of interest by averaging over 500 × 6 = 3000 samples. For more details about the simulations see Supplement section S8.

### Resources

Documented code for the simulations and numerical solutions of the model, as well as the scripts used to produce all figures can be downloaded at https://github.com/jjberg2/msdbPaperCode.

## Results

### The population dynamics and genetic architecture at individual sites

#### The dynamics at a site

The dynamics at a site can be described in terms of the first two moments of change in allele frequency in a single generation (Ewens 2004, Ch. 4). We calculate the moments by averaging the fitness of the three genotypes over genetic backgrounds and plugging these averages into the standard equation for the change in allele frequency at a single locus (Supplement section 2.1). We find that the expected change in frequency of an allele at frequency *x* that increases liability by *a* is well approximated by

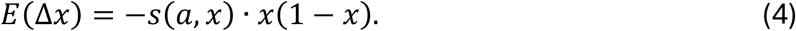

The selection coefficient *s*(*a, x*) takes an intuitive form: it equals the fitness cost of the disease, *C*, multiplied by the allele’s effect on disease risk, *δ*_*R*_(*a, x*), namely:

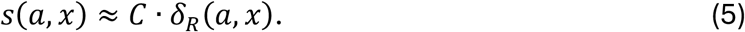

*δ*_*R*_ (*a, x*) is defined as the expected increase in individual disease risk caused by substituting a random liability-decreasing allele at this site by a liability-increasing one. As an aside, we note that the allele’s effect on risk equals its (absolute) “population attributable risk” in epidemiology and can be translated into its odds-ratio estimated in case-control GWAS (see Supplement section 2.5). The second moment of change in allele frequency is well approximated by the standard drift term

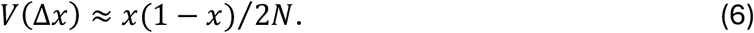

To complete the description of these dynamics, we require the functional form of the risk effect *δ*_*R*_(*a, x*). In Supplement section 3, we show that the risk effect is well approximated by the area under the liability distribution that is pushed over the liability threshold when the distribution is shifted by *a* on the liability scale (Figure 2). We denote the probability density at liability *Z* by *f*(*Z*) and the probability that liability exceeds liability *Z* by *F*(*Z*). In these terms,

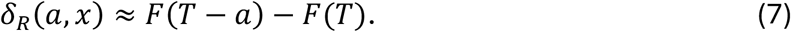

**Figure 2.**
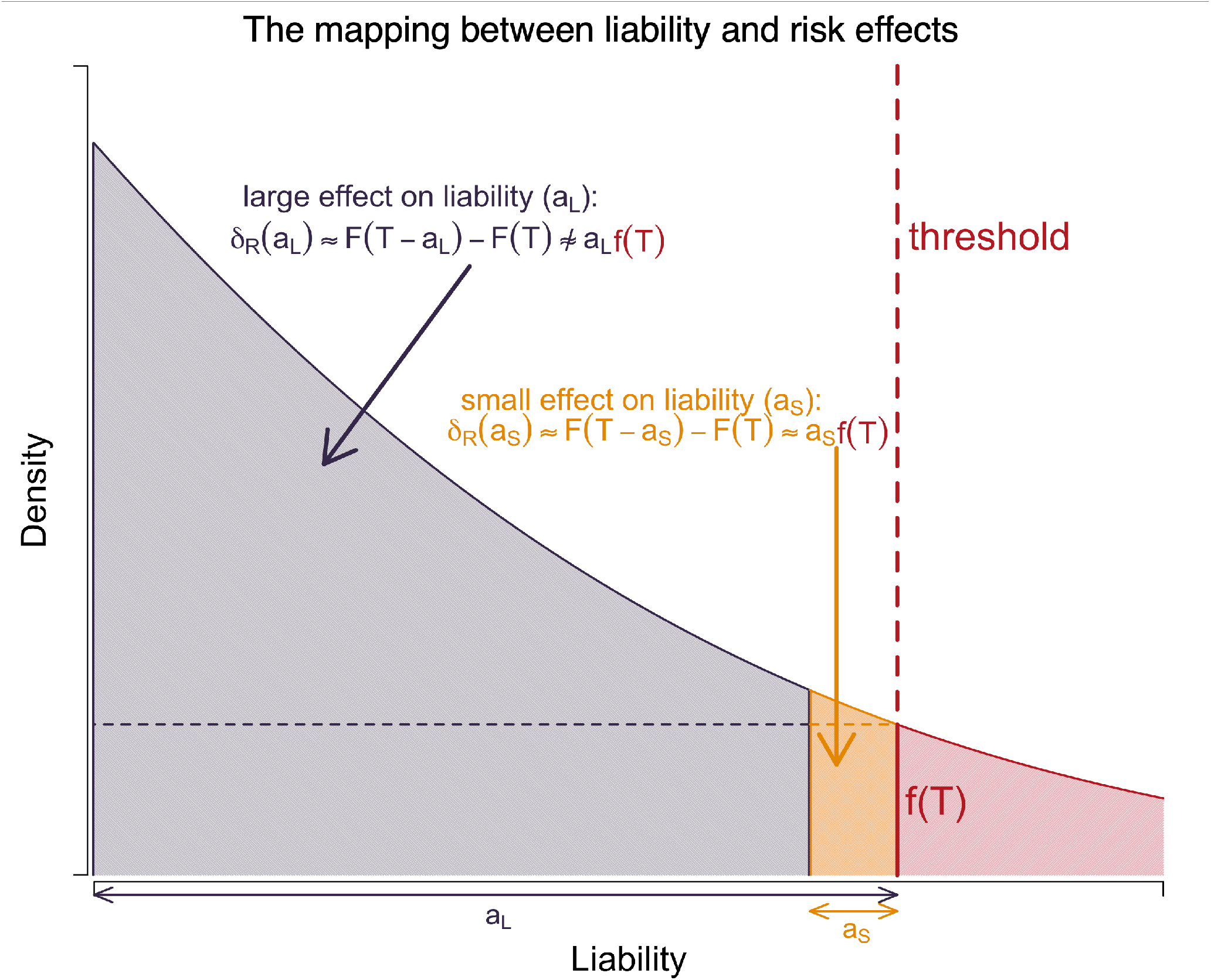
The mapping between liability and risk effects. For small effect sites, illustrated in yellow, the risk effect is approximately equal to the product of the liability effect and the threshold density. For large effect sites, this linear approximation is inaccurate, and the risk effect must be approximated in terms of the difference in the areas in the tails of the liability distribution.

We can therefore assume that the risk effect and selection coefficient do not vary with allele frequency and adjust our notation to *δ*_*R*_(*a*) and *s*(*a*), dropping the dependence on *x*.

The dependence of an allele’s risk effect on its liability effect can be divided into two cases (Figure 2). When the allele’s effect on liability (*a*) is sufficiently small such that the corresponding shift to the liability distribution has a negligible effect on the probability density near the threshold (*T*), we can approximate the allele’s effect on risk by the area of the rectangle with width *a* and height *f*(*T*), i.e.,

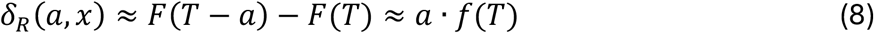

(Milkman 1978; Kimura and Crow 1978). An allele is small in this sense if its effect is substantially smaller than the width of the phenotypic distribution, i.e., 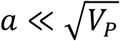 (see Supplement section 2.3 for further analysis).

In turn, an allele is large if its effect on liability is comparable to or greater than the width of the phenotypic distribution, i.e.,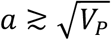 . When the effect of alleles on liability increases from small to large, the dependence of their risk effect *δ*_*R*_ on their liability effect (*a*) becomes superlinear (Figure 2; but see Figure S1). In Supplement section 3.4, we show that for the kinds of diseases that we consider–that have a substantial fitness cost and are not exceedingly rare–alleles become strongly selected (i.e., 2*Ns*(*a*) ≈ 2*NCδ*_*R*_(*a*) ≫ 1) before the superlinear dependence kicks in. This finding implies that we can divide alleles into three categories: small (in the sense of Eq. 8) and weakly selected (i.e., 2*Ns*(*a*) ≅ 2*NCδ*_*R*_(*a*) ≲ 1), small and strongly selected, and large and strongly selected.

Next, we consider the genetic architecture at mutation-selection-drift balance (MSDB). To this end, we only care about alleles’ selection coefficients (rather than their effects on liability). The insensitivity of alleles’ risk effect to their frequency allows us to treat their selection coefficients as constant, which, in turn, allows us to apply standard approximations to solve for quantities of interest throughout the range allele effect sizes.

#### The fixed state

At MSDB, fixations are at detailed balance: for a given effect size, the rates of fixation of liability-increasing and -decreasing alleles at sites are equal, i.e., they balance each other out (Iwasa 1988; Sella and Hirsh 2005). The proportions of sites with effect *a* fixed for the risk increasing and decreasing alleles, *p*_+_(*a*) and *p*_−_(*a*) = 1 − *p*_+_ (*a*) respectively, therefore satisfy

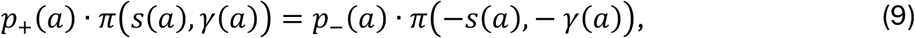

where *π*(*s, γ*) ≈ 2*s*/(1 − *e*^−2*γ*^) is the fixation probability of a mutation with selection coefficient *s* and scaled selection coefficient *γ* = 2*Ns* that arises at frequency 1/2*N* (Crow and Kimura 1970). We solve for the fixed state and represent the solution in terms of the bias toward risk-decreasing alleles

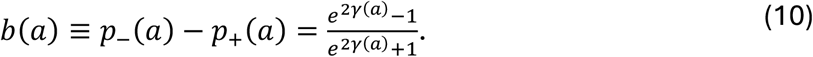

In these terms, *p*_+_(*a*) = (1 − *b*(*a*))/2 and *p*_−_(*a*) = (1 + *b*(*a*))/2. The expected bias satisfies 1 > *b*(*a*) > 0, because selection always favors the risk-decreasing allele.

The fixed state exhibits the three standard selection regimes (Figure 3A). When selection is extremely weak and drift dominates, sites are equally likely to be fixed for the risk-increasing and -decreasing alleles, i.e., *b*(*a*) ≈ 0. This effectively neutral regime occurs when *γ*(*a*) ≈ 2*NC*δ_*R*_(*a*) ≪ 1. At the other extreme, when selection is strong and dominates over drift, sites are always fixed for the risk-decreasing allele, i.e., *b*(*a*) ≈ 1. This strong selection regime occurs when *γ*(*a*) ≈ 2*NCδ*_*R*_(*a*) ≫ 1. In the weak selection regime, when *γ*(*a*) ≈ 2*NCδ*_*R*_(*a*)∼1, the fixed state transitions between the effectively neutral and strongly selected extremes, with the bias *b*(*a*) increasing between 0 and 1 as selection becomes stronger.

**Figure 3.**
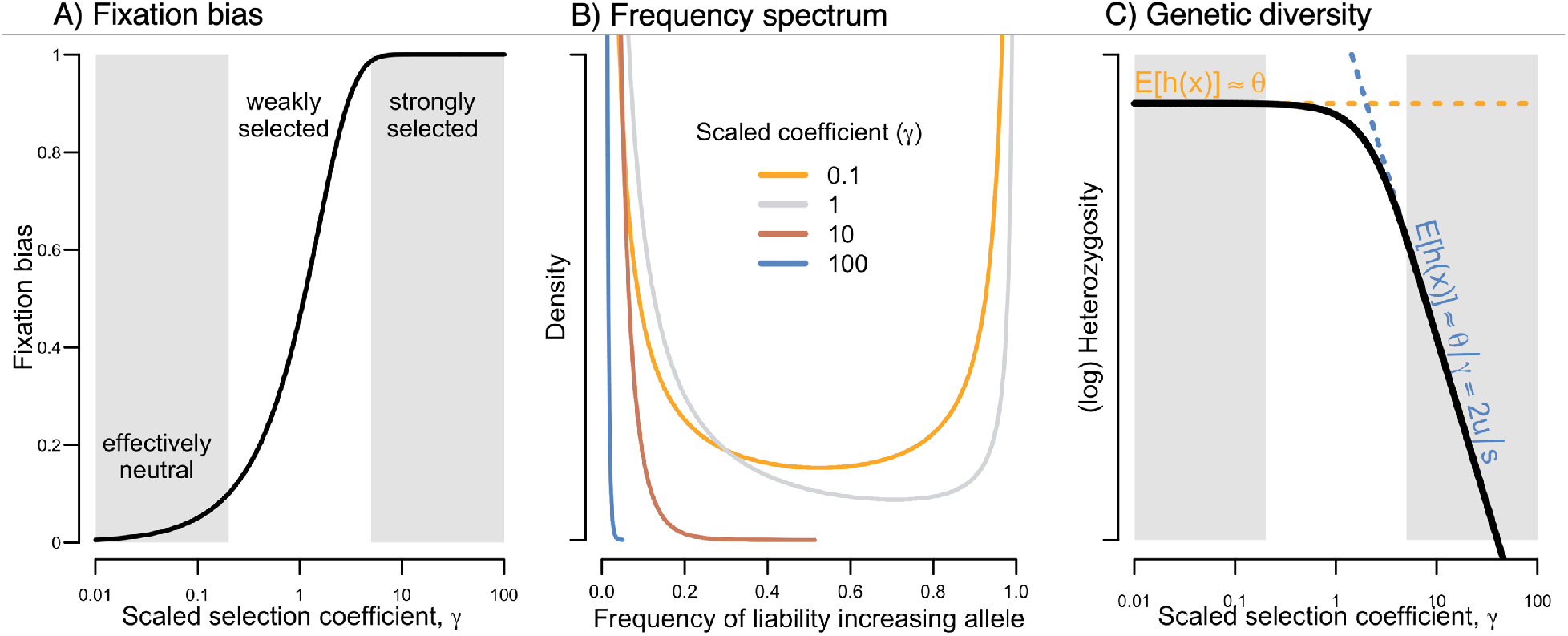
The genetic architecture at MSDB. A) The fixation bias as a function of the scaled selection coefficient at a site. B) The site frequency spectrum. C) The expected genetic diversity as a function of the scaled selection coefficient.

With the fixed state biased toward risk-decreasing alleles (at all but effectively neutral sites), mutation is biased toward risk-increasing alleles. Like in the classic model of simple (Mendelian) diseases, this mutational asymmetry arises from the dynamics of the model rather than from assumptions about mutation (namely, we assumed symmetric mutation between risk-increasing and -decreasing alleles).

#### Segregating sites

Figure 3B shows the frequency distribution of segregating, disease-increasing alleles at MSDB for several values of the population-scaled selection coefficient (see Supplement section 3.1 for derivation). With the fixed state biased toward alleles that decrease risk, derived, segregating alleles tend to increase risk. Even neutral derived alleles, let alone derived alleles that increase risk, segregate at lower frequencies than ancestral ones. These effects explain why risk-increasing alleles tend to segregate below frequency ½, and why this bias is stronger when selection is stronger. The predicted asymmetry between the frequencies of alleles that increase and decrease risk can be tested using data from GWAS (see Discussion and Koch et al., 2024).

Next, we consider diversity levels. We can calculate the expected diversity levels using the diffusion approximation (Crow and Kimura 1970; Ewens 2004). A variant with allele frequency *x* contributes *h*(*x*) = 2*x*(1 − *x*) to heterozygosity. To calculate the expectation per site, we multiply the rate at which mutations arise by the expected total contribution of an individual mutation during its sojourn in the population. Namely,

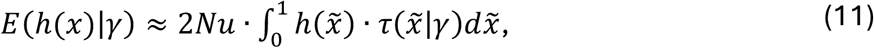

with the sojourn time

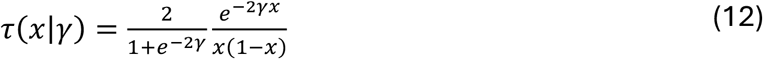

defined such that *τ*(*x*|*γ*)*dx* is the expected number of generations that an allele with scaled selection coefficient *γ* spends between frequencies *x* and *x* + *dx*. In this way, we find that the expected heterozygosity per site

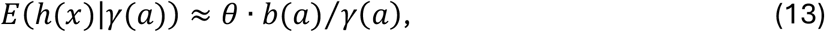

where *θ* = 4*Nu* and *b*(*a*) is the fixed bias.

Diversity levels also exhibit the three standard selection regimes (Fig. 3C). In the effectively neutral regime (i.e., when *γ* ≪ 1), heterozygosity is well approximated by the neutral expectation *θ*. In the strong selection regime (i.e., *γ* ≫ 1), sites are always fixed for the risk decreasing allele, implying that *b*(*a*) ≈ 1 and that the derived, risk increasing allele segregates at low frequency, i.e., *x* ≪ 1, so

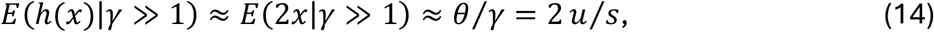

which aligns with the classic expectation under mutation-selection balance. In the weak selection regime (i.e., when *γ*∼1), diversity levels transition between these two extremes.

#### Contribution to variance

The last facet of architecture that we consider here is the contribution to additive variance in liability. Estimates of this contribution are used to assess how much of the heritable variance in disease risk arises from variants of small and large effects (see Discussion). The total genetic variance in liability will become important when we consider disease prevalence.

We can calculate the expected contribution to variance based on the expected heterozygosity (Eq. 13). A variant with allele frequency *x* and liability effect *a* contributes *v*(*a, x*) = 2*a*^2^*x*(1 − *x*) = *a*^2^ · *h*(*x*) to additive variance in liability. Therefore, the expected contribution per site

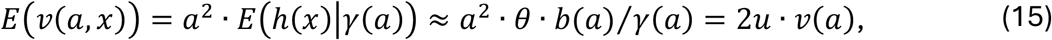

where

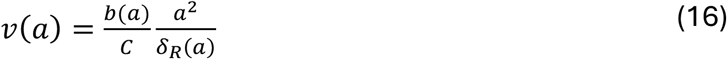

is the contribution per unit diploid mutation rate.

We can calculate the total additive variance in liability by summing over sites and integrating over the distribution of effect sizes. Namely,

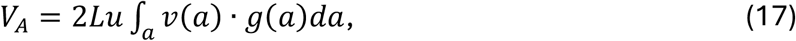

where *L* is the number of sites and *g* is the distribution of effect sizes. If we assume that all effect sizes are small, then *δ*_*R*_(*a*) ≈ *a* · *f*(*T*) (Eq. 8) and *v*(*a*) ≈ *b*(*a*) · *a*/Y*C* · *f*(*T*)). In this case, the total variance is well approximated by

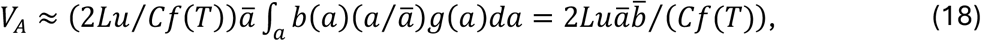

where 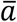 is the mean effect size and 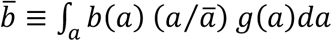 is the mean fixation bias, with sites weighted by their effect sizes.

In Supplement section S4.3, we investigate how the contributions to liability- and risk-scale variance vary with variant effect sizes. We show that in our model, these variances do not exhibit the same kind of asymptotic ‘flattening’ found in models of stabilizing selection (see Discussion and Simons et al. 2018; O’Connor et al. 2019). We also describe how some simple summaries of asymmetry in the architecture at individual sites can be calculated (Supplement section S4.4; Figure S12 and S13). Like the summaries considered here, all these summaries be described in terms of *a, θ* and *γ*(*a*).

### The mapping between liability effects and selection coefficients

#### A phenotypic perspective on MSDB

At MSDB, mutation is biased toward risk-increasing alleles, causing a mutational increase in mean genetic liability each generation; we denote it by 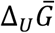. The mutational bias is balanced by an equal but opposing selection response that we denote by 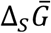. Thus, at MSDB, 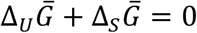.

Here we focus on the mutational bias. We first approximate the mutational bias based on the fixed state. In this approximation, all risk-increasing mutations occur at sites fixed for the risk-decreasing allele, and vice versa. As above, we denote the proportions of sites with effect *a* fixed for the risk-increasing and -decreasing alleles at MSDB by *p*_+_(*a*) and *p*_−_ (*a*) = 1 − *p*_+_(*a*), respectively. We then approximate the mutational bias as

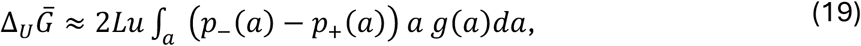

where *g*(*a*) is the distribution of liability effects across sites. Recalling that the fixation bias *b*(*a*) ≡ *p*_−_(*a*) − *p*_+_ (*a*), we find that

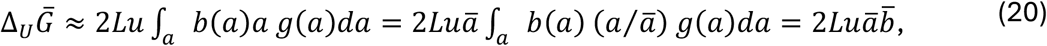

where 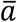 is the mean effect size and 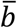 is the mean fixation bias, weighted by effect sizes.

We can also express the mutational bias in terms of the mean genetic liability 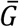. In the fixed state approximation, the mean liability is

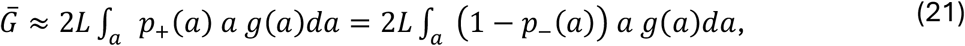

where we have set the range of genetic liability scale between 0 and 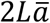, with low liability alleles contributing 0. From Eqs. 19-21, we find that

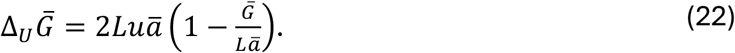

In Supplementary section 5, we show that this equation is exact when we relax the fixed state approximation and account for segregating genetic variation.

Under our modeling assumptions, the distance between the mean genetic liability 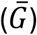 and the liability threshold (*T*) is tiny relative to the scale of possible genetic liability 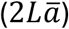, i.e.,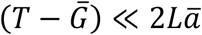. To understand why, we consider the case without an environmental contribution to liability. (An environmental contribution only reduces the distance between the mean genetic liability and the threshold, because it reduces the efficacy of selection on individual sites.) Without an environmental contribution, our assumption of a low mutation rate per site (*θ* = 4*Nu* ≪ 1) implies that the scale of variation in liability among individuals is much smaller than the scale of possible genetic liabilities (i.e.,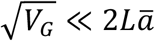). In turn, our assumption that the disease prevalence is not exceedingly small requires the distance between the mean genetic liability and the threshold to be on the scale of the genetic variation (i.e.,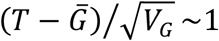). It therefore follows that 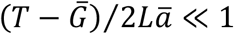.

This condition allows us to approximate the mutational bias in terms of the position of the threshold. Specifically, given that 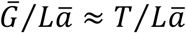, we find that

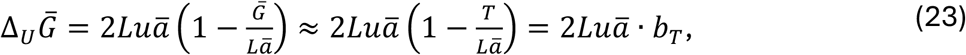

where 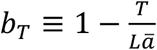 measures the position of the threshold relative to the middle of the liability scale; we henceforth refer to it as the threshold bias. Eq. 23 shows that the threshold bias determines the mutational bias. Moreover, comparing Eqs. 20 and 23, we find that

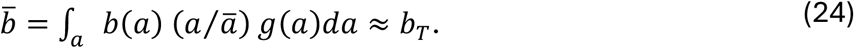

Thus, the threshold bias also determines the mean fixed bias at MSDB, which reflects the strength of selection acting on individual sites (Eq. 10).

#### Selection at sites with small effects

Equation 24 also tells us how the strength of selection on sites relates to their effects on liability, so long as these effects are small. To understand how, we first consider a simple case in which all sites have the same liability effect size *a* and therefore the same scaled selection coefficient *γ*(*a*). When we express the fixation bias in terms of the scaled selection coefficient (Eq. 10), Eq. 24 becomes

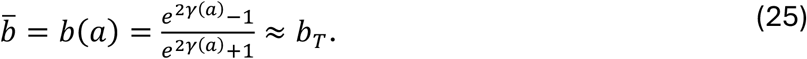

Solving for the scaled selection coefficient, we find that

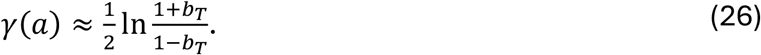

Thus, the position of liability threshold *b*_*T*_ determines the scaled selection coefficient at MSDB *γ*(*a*) (Fig. 4A). Eq. 26 applies so long as the threshold is not extremely close to 0 (specifically, *T*/(*La*) = 1 – *b*_*T*_ ≫ 1/*L*), such that some sites are fixed for the liability-increasing allele. For this to be the case selection cannot be too strong, implying that our small effect size approximation (Eq. 8) applies and that *γ*(*a*) ≈ 2*NCδ*_*R*_(*a*) ≈ (2*NCa*) · *f*(*T*). Consequently, the threshold density at MSDB is

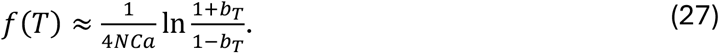

**Figure 4.**
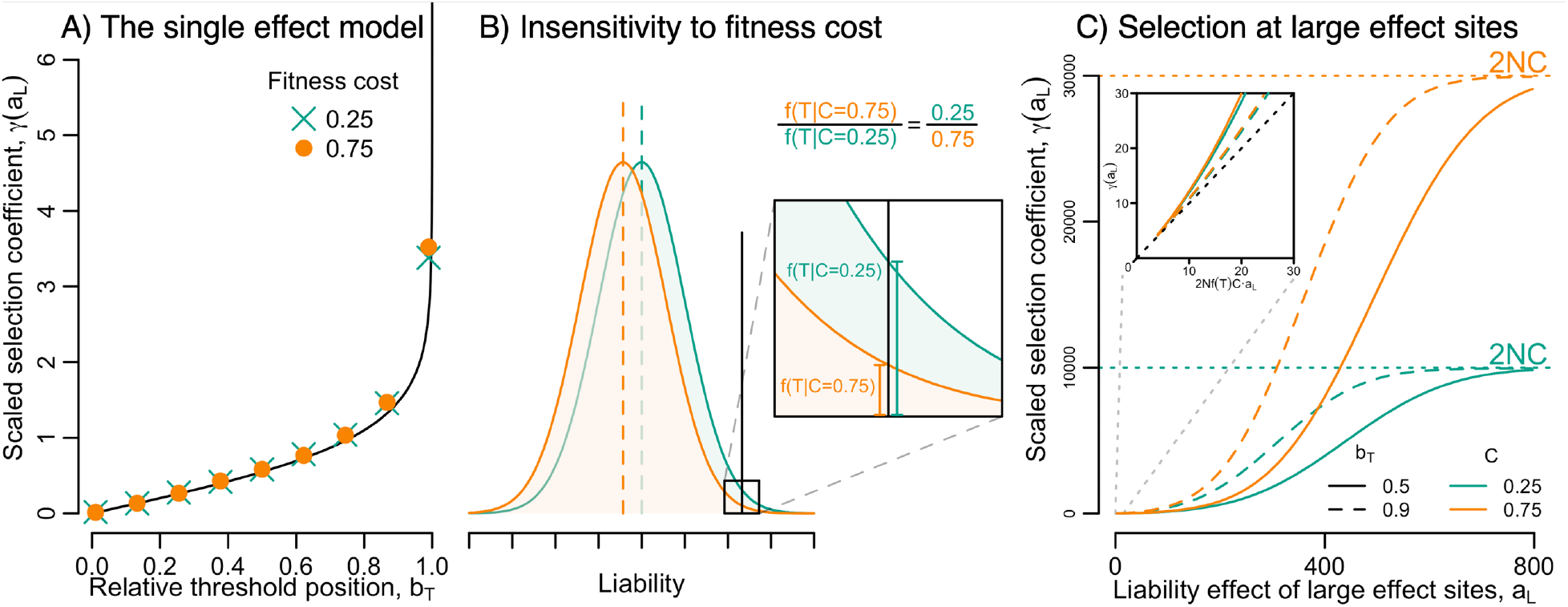
The mapping between liability and fitness effects. A) The threshold position determines the scaled selection coefficients in the model with a single effect size. The solid line depicts the analytic approximation (Eq. 6) and the circles and crosses depict averages in simulations with specified fitness costs (see Model and Supplement section S8 for details). Scaled selection coefficients in simulations were estimated by computing the average of the risk effect, *δ*_*R*_(*a*), over many sites and generations and multiplying by 2*NC*. B) Selection on small effect sites is insensitive to the cost of the disease. When the fitness cost, *C*, increases, mean liability is pushed down to reduce the threshold density, *f*(*T*), such that 2*NCf*(*T*) and thus scaled selection coefficients at small effect sites remain invariant. C) The mapping between liability and scaled selection effects at large effect sites. The results shown are based on numerically solving the model with two effect sizes (see Supplement section S7 for details), setting *N* = 20,000, *L* = 1.5 × 10^7^, and *u* = 10^−8^ per site per generation, with fractions *p*_*S*_ = 0.9995 of small effect sites and *p*_*l*_ = 0.0005 of large effect sites, varying *a*_*l*_ with *a*_*S*_ = 1, and using the specified values of *b*_*T*_ and *C*.

Thus, given the compound evolutionary parameter 4*NCa*, the required strength of selection is attained by adjusting the population’s liability distribution, such that the threshold density *f*(*T*) satisfies Eq. 27.

Next, we consider the general case with a distribution of effect sizes. In this case, when we express Eq. 24 in terms of scaled selection coefficients, we find that

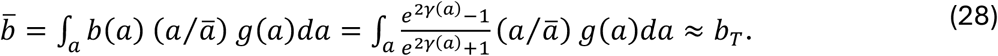

If we assume the small effect approximation for *γ*(*a*), the equation becomes

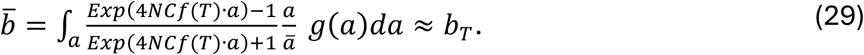

We can solve this equation numerically using a line search for the threshold density *f*(*T*) given the distribution *g*, 4*NC*, and *b*_T_. Importantly, given the *f*(*T*) that solves Eq. 29, *γ*(*a*) ≈ (2*NCa*) · *f*(*T*) solves Eq. 28.

The solution for the threshold density divides sites into three kinds. On the high end of effect sizes, sites are strongly selected, with (2*NCa*) · *f*(*T*) ≫ 1, and thus fixed for the liability decreasing alleles; these sites all contribute *b* ≈ 1 to attaining 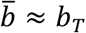. On the lower end of effect sizes, sites are effectively neutral, with (2*NCa*) · *f*(*T*) ≪ 1, and thus equally likely to be fixed for the liability-increasing and -decreasing alleles; these sites all contribute *b* ≈ 0 to attaining 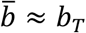. In between these ends, sites are weakly selected, with (2*NCa*) · *f*(*T*)∼1 and their fixed state is highly sensitive to the threshold density, with the fixed bias *b* ranging between 0 and 1.

If the distribution of liability effects *g* is highly concentrated around 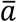, the solution resembles the case with a single effect size. MSDB is attained by adjusting the threshold density such that most sites—with effects near 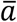—are weakly selected, and 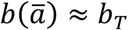. In the other extreme, with liability effects thinly distributed over several orders of magnitude, only a small proportion of sites would fall in the intermediate, weakly selected range of effect sizes. In this case, the threshold density at MSDB divides the range of effect sizes such that the fixed bias from strongly selected sites matches the threshold bias, i.e., such that 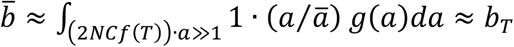. In Supplementary section 6, we investigate how variation in the distribution of liability effects, *g*, and relative position of the liability threshold, *b*_*T*_, would affect the solution of Eq. 29, assuming most sites have small effects.

Here we focus on what the solution of Eq. 29 tells us about the mapping between liability effects and selection coefficients. As we already know, the approximation *γ*(*a*) ≈ (2*NCa*) · *f*(*T*) breaks down when sites are sufficiently large (see, e.g., Fig. 2). This does not affect the solution to Eq. 29 because selection at these sites is sufficiently strong (i.e., *γ*(*a*) ≫ 1) to maximize their fixed bias (i.e., *b*(*a*) ≈ 1) regardless of the exact form relating their liability effect sizes and selection coefficients. This reasoning clarifies that Equations 29 is insensitive to, and uninformative about, the strength of selection acting on sites with sufficiently large effects.

In contrast, the strength of selection acting on sites with small effects follows from Equation 29. When effect sizes are sufficiently small, the strength of selection is well approximated by *γ*(*a*) ≈ (2*NCa*) · *f*(*T*). As we already noted, for diseases with a substantial fitness cost that are not exceedingly rare, sites with such small effect liabilities range from being effectively neutral to being strongly selected. Selection on sites in this range is determined by the relative position of the liability threshold, *b*_*T*_, and the distribution of effects, *g*, where the mapping between effect sizes and scaled selection coefficients is attained by adjusting the threshold density, *f*(*T*).

#### The genetic architecture at sites with small effects

Our results imply that scaled section coefficients at sites with small effects are insensitive to the fitness cost, *C*, environmental variance, *V*_*E*_, and population size, *N*. Figure 4B illustrates the effects of an increase in fitness cost. In this case, the mean genetic liability 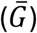 is pushed farther below the threshold to reduce the threshold density (*f*(*T*)), leaving 4*NCf*(*T*) unchanged, and maintaining the same mapping between liability effect sizes and scaled selection coefficients. An increase in population size or decrease in environmental variance act similarly, although they also affect the total variance in liability. Because the disease is highly polygenic, changes in mean genetic liability are achieved by tiny changes to the fixed state across sites, with negligibly small effects on the scaled selection coefficients.

As we described earlier, the architecture at sites depends only on *a, θ* and *γ*(*a*). For sites with small effects, the dependence on *γ* translates into a dependence on the threshold bias, *b*_*T*_, and the distribution of liability effects, *g*. In turn, the architecture at sites with small effects is insensitive to the fitness cost of the disease and the environmental variance, while the population size affects only the total number of segregating sites, but not the distribution of their allele frequencies.

#### Selection on and genetic architecture of sites with large effects

Large effect sites span a wide range of liability effect sizes, and the factors that determine the strength of selection acting on them vary within this range. At the lower end of this range, liability effect sizes are just above those of strongly selected, small effect sites. Near this boundary, scaled selection coefficients are still strongly affected by the threshold density *f*(*T*), as well as by derivatives of the liability distribution at the threshold (see, e.g., Figure 2). We would therefore expect selection on such large effect sites to resemble selection on strongly selected, small effect sites in being affected by the relative position of the threshold, *b*_*T*_, and distribution of selection effects, *g*.

At the other end, we have sites with effect sizes that are large enough for a single copy of the risk-increasing allele to almost always cause the disease (e.g., when 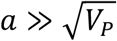); these alleles are dominant, with full or nearly full penetrance. At this end, *δ*_*R*_ (*a*) ≈ 1, *s* ≈ *C, γ* ≈ 2*NC*, and the expected frequency of the risk-increasing allele at a site is describe by classic MSDB, with *E*(*x*) ≈ *u*/*C*. Thus, in contrast to sites with large effect sizes at the lower end (and to sites with small effects), the selection acting on them and their genetic architecture are determined by the fitness cost of the disease and are insensitive to the relative position of the liability threshold, *b*_*T*_.

Figure 4C shows the mapping between effect size and scaled selection coefficients for a large effect site, in a model with two effect sizes. When the large effect size increases, the dependence of the strength of selection on model parameters varies gradually between the two behaviors that we described.

### Disease prevalence

Lastly, we consider the disease prevalence at MSDB. The prevalence is equal to the area under the tail of the liability distribution that lies beyond the threshold, so calculating it requires knowledge of the shape of this distribution. The liability distribution is often assumed to be Normal (see Discussion). This assumption seems sensible when the disease is highly polygenic and genetic contributions are small, such that an individual’s liability arises from many small effect genetic contributions and a normally distributed environmental contribution.

We begin by considering this case and assuming normality, which allows us to calculate the prevalence based on the density of the liability distribution at the threshold. Given the threshold density on the standard scale, *φ*_*T*_, the prevalence is given by

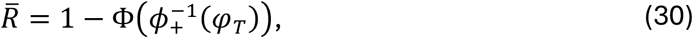

where 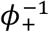 is the positively defined inverse of the standard Normal PDF, and Φ is the standard Normal CDF. As we assume that the disease is not exceedingly common or exceedingly rare (e.g., 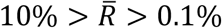), the dependence on the threshold density is approximately linear, with

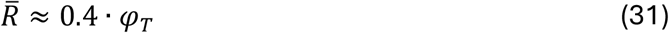

(Fig. S2).

We can rely on our small effect approximations to calculate the threshold density on the standard scale. The standardized threshold density

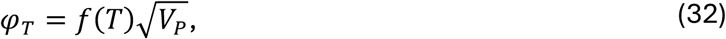

where *V*_*P*_ is the variance in liability. Noting that in the small effect approximation 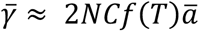, that the heritability in liability *h*^2^ = *V*_*A*_/*V*_*P*_ and relying on our approximation for *V*_*A*_ (Eq. 18), we find that

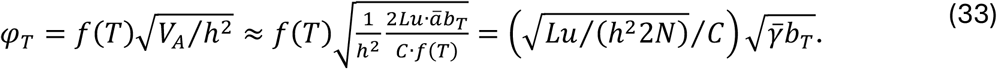

Under our assumption that effect sizes are small, the mean scaled selection coefficient 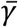 and thus the term 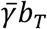 are fully determined by the threshold bias, *b*_*T*_, and distribution of effect sizes, *g*. In the single effect case, we can substitute the explicit expression for 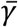 (Eq. 26) to find that the standard threshold density

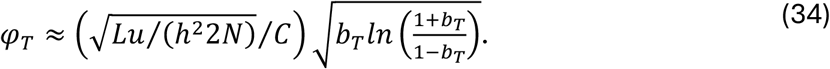

Figure 5A shows how the prevalence in the single effect size model increases with *b*_*T*_ for several possible values of the compound parameter 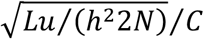. As an illustration, we consider parameter values typical of humans, i.e., *N* = 2 · 10^4^ and *u* = 10^−8^ per site per mgeneration, a heritability *h*^2^ = 1/2, and a cost *C* = 0.1. We vary the target size within the wide range estimated for complex, quantitative traits, e.g., *L* = 1.5 · 10^6^— 1.5 · 10^8^ (Simons et al. 2022). For these parameters, we find that a disease prevalence of 1% or greater at MSDB is attainable only if the target size or the threshold bias is relatively large.

**Figure 5.**
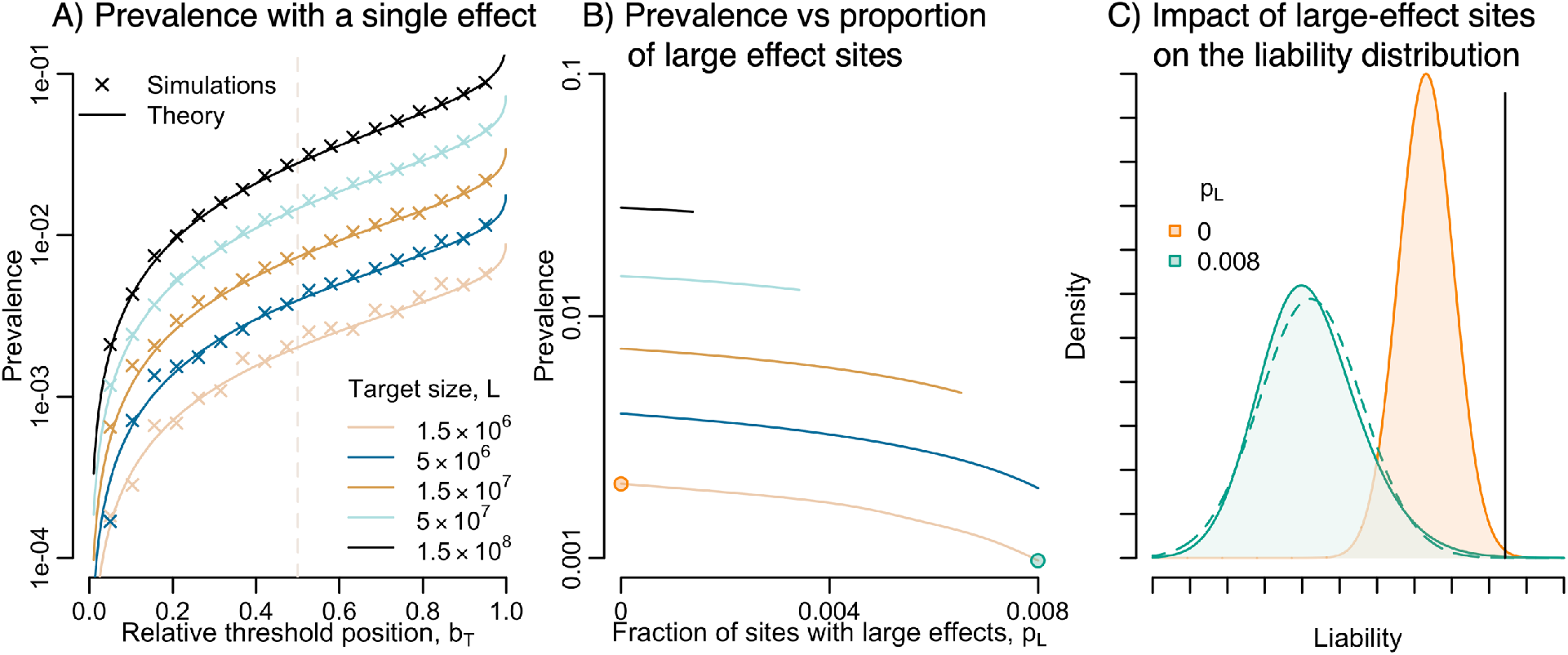
Disease prevalence at MSDB. A) Prevalence versus threshold bias in the model with a single effect size. Analytic results are based on Eqs. 30 and 34; for simulations details see Methods and Supplement section S8. The results correspond to setting *N* = 2 · 10^4^, *u* = 10^−8^ per site per generation, *h*^2^ = 1/2, and *C* = 0.1, and varying the target size *L* and threshold bias *b*_*T*_ as indicated. B) Prevalence versus the fraction of sites with large effect sites in the model with two effect sizes. The model was solved as described in the text (see Supplement section S7 for details). The parameters are as in A, with *b*_*T*_ = 1/2 and *a*_*l*_/*a*_*S*_ = 100. See Figure S3 for other choices of *b*_*T*_ and *a*_*l*_/*a*_*S*_. C) The impact of large effect sites on the liability distribution. The two distributions shown correspond to the parameter values highlighted in panel B, with and without large effect sites. The dotted outline shows a Normal distribution with the same mean and variance as the Poisson convolution with large effects.

Next, we consider how sites with large effects impact prevalence. To this end, we study a model with two effect sizes: a fraction *p*_*S*_ of sites have a small effect size *a*_*S*_ and a fraction *p*_*l*_ of sites have a large effect size *a*_*l*_, where *p*_*S*_ + *p*_*l*_ = 1. In this parametrization, the boundary case with *p*_*S*_ = 1 and *p*_*l*_ = 0 corresponds to the single effect model that we considered before, and changes in prevalence when we move away from this boundary by increasing *p*_*l*_ reflect the effect of increasing the proportion of large effects.

If we plausibly assume that individuals carry only a few large effect risk-increasing alleles, then we no longer expect the liability distribution to be Normal. Variation among individuals in the number of these alleles introduces a fat tail of individuals with higher liabilities and thus a skewed liability distribution. In the case with two effect sizes, the number of large effect, risk-increasing alleles follows a Poisson distribution with mean 2*Lup*_*l*_/*s*(*a*_*l*_) (Felsenstein 1974), where we assume that the mean 2*Lup*_*l*_/(*s*(*a*_*l*_)) ≤ 3 so that an individual carries only at most a few of these alleles.

We therefore model the liability distribution at MSDB as arising from three components: (1) a large effect genetic liability distribution that arises from the Poisson distributed number of large effect risk-increasing alleles, (2) a normally distributed genetic liability arising from small effect sites, and (3) a normally distributed environmental liability. The liability of an individual in the population is randomly sampled from the contributions to liability arising from each of these components. The liability distribution is therefore the convolution of these three distributions.

We solve this model numerically (for details see Supplement section S7). We assume that all large effect sites are fixed for the low liability allele and derive the threshold density, *f*(*T*), by requiring that the fixed bias from the small and large effect sites combined matches the threshold position, *b*_*T*_. Given the threshold density, we solve for the liability distribution arising from small effect sites. We then solve for the selection coefficient of large effect sites, *s*(*a*_*l*_), by requiring that the convolution of the Normal and Poisson components of the liability distribution match the threshold density, *f*(*T*).

Figure 5B shows the prevalence as a function of the proportion of large effect sites. Here, we set *a*_*l*_/*a*_*S*_ = 100 and *b*_*T*_ = 1/2 but in Figure S3, we explore other choices and obtain similar qualitative results. All other model parameters are set to the same values that we used for the case with a single effect size (as in Fig 5A). We vary the proportion of large effect sites between *p*_*l*_ = 0, corresponding to the case without large effects, and *p*_*l*_ such that (2*Lup*_*l*_/*s*(*a*_*l*_)) = 3. We validated our numerical solution against simulations (Fig. S7).

Increasing the proportion of large effect sites affects the liability distribution in three ways (Figs. 5C). First, it increases the variance in liability, because large effect sites contribute much more variance than small effect sites (Supplement section S4.3). Second, it introduces a right skew in liability due to variation in the number of large effect risk-increasing alleles. Third, it decreases the threshold density *f*(*T*), because, with large effect sites all fixed for the risk-decreasing alleles, the fixed bias and thus the selection at small effect sites is weaker. We would expect the first two effects to increase the prevalence and the third one to decrease it. For the parameter values that we consider in Fig. 5B, the combination of these effects leads to a reduction in prevalence when the fraction of large effect sites increases. In Fig. S3-S6, we consider how the prevalence changes with increasing fraction of large effect sites for other choices of *b*_*T*_ and *a*_*l*_/*a*_*S*_. In general, we find that when *b*_*T*_ is closer to zero, increasing the fraction of large effect sites tends to lead to a decrease in prevalence, whereas when it is closer to one, it tends to lead to an increase.

## Discussion

We introduced an evolutionary model of complex disease susceptibility, in which a variant’s effect on disease risk follows from the liability threshold model, and a variant’s effect on fitness follows from its effect on disease risk. The model can be viewed as a generalization of the classic MSDB model of Mendelian diseases and is also closely related to models used to study genetic load (see Introduction). We solved the model for the genetic architecture and prevalence of the disease at MSDB.

Mutation-selection-drift balance in this model can be understood as a ‘mean field’ equilibrium. Selection on sites with a given effect on liability is determined by the phenotypic distribution (the ‘field’), and the phenotypic distribution arises from the aggregate behavior at all sites (the ‘mean’). The density of the phenotypic distribution at the liability threshold is determined by matching the population’s fixed state with the position of the threshold on the liability scale (given the distribution of liability effect sizes). The threshold density divides liability effect sizes into ‘small’ and ‘large’, which differ in their mapping onto the effects on disease risk and fitness. For sites with small effects on the liability scale, the effects on disease risk and fitness are proportional to their liability effects and to the threshold density. For sites with large effects, the effects on disease risk and fitness depend on non-linear cumulants of the liability distribution and the fitness cost of the disease. The selection acting on sites shapes their genetic architecture, which together with environmental effects on liability, determines disease prevalence. We mapped out these relationships and their main implications for observable quantities and solved the model explicitly for simple distributions of effect sizes.

With these predictions in hand, we can ask whether our model fits what is known about the genetic architecture of complex disease susceptibility in humans. Figure 6 shows examples of the “smile” architecture of GWAS hits typical of complex diseases (Koch et al. 2024). These hits were ascertained in GWAS based on genotyping and imputation and therefore include common variants under weak and moderate selection but not variants under very strong selection. The “smile” architecture reflects selection in that variants with larger risk effects segregate at low minor allele frequencies (i.e., risk allele frequencies near 0 or 1). This is not the signature of selection expected under our model. Instead, we predict that if selection acted on variants due to their effects on disease risk, the architecture should be asymmetric, with a depletion of major alleles increasing risk, rather than (approximately) symmetric between minor and major risk-increasing alleles (compare Figs. 3B and 6).

**Figure 6.**
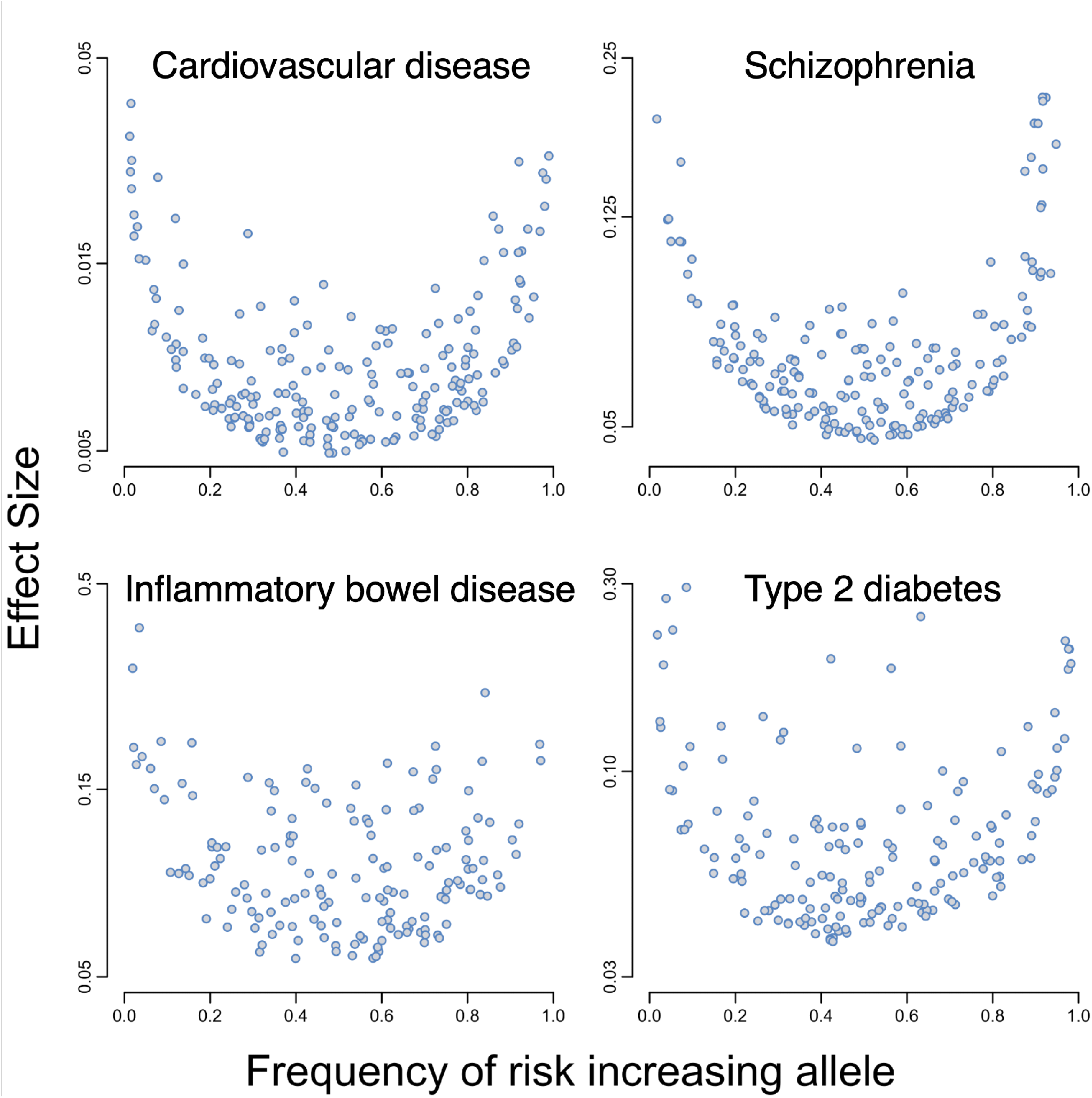
The “smile” architecture of complex diseases in humans. Points correspond to approximately independent genome-wide significant hits. The data was taken from Dönertaş et al (2021), Trubetskoy et al. (2022), Liu et al. (2015), and Spracklen et al. (2020). See Supplement section S9 for data processing details.

Our results suggest that selection on variants due to their effects on disease risk should be stronger for diseases with greater fitness costs and higher prevalences (see, e.g., Eqs. 5 and 34). The departure from our predictions might seem all the more surprising then, given that complex diseases often entail substantial fitness costs and are quite common in contemporary human populations. Schizophrenia, for example, has a prevalence of 0.5-1% (Jablensky 2000; Saha et al. 2005; Chan et al. 2015; Simeone et al. 2015), affects several fitness components and has been estimated to reduce fertility by up to 75% (Haukka, Suvisaari, and Lönnqvist 2003; Laursen and Munk-Olsen 2010; Power et al. 2013). Other diseases, such as type 2 diabetes, with a global prevalence of 6% (Ong et al. 2023) and multiple sclerosis, with a prevalence of ∼0.3% in the United States (Nelson et al. 2019), are associated with a substantial increase in mortality rates and are plausibly associated with substantial reductions in fitness (Scalfari et al. 2013; Graves et al. 2023; Emerging Risk Factors Collaboration 2023). How then might we make sense of the fact that the architecture of common variation affecting complex disease susceptibility does not reflect selection against these diseases?

One possibility is that complex diseases that are common in contemporary human populations substantially increased in prevalence with very recent changes in environment. Examples plausibly include type 2 diabetes and other diseases associated with obesity (Dai et al. 2020; Teng et al. 2022; Ong et al. 2023), asthma (Eder, Ege, and Mutius 2006), and autism (Atladóttir et al. 2007; Weintraub 2011; Hansen, Schendel, and Parner 2015). More generally, we know little about the fitness cost and prevalence of human diseases more than a century back.

Persistent selection over molecular evolutionary time scales would be needed to attain the fixed bias that shapes the genetic architecture at MSDB in our model. These time scales vary with the scaled selection coefficients affecting sites. As an illustration, given the contemporary mutation rate in humans and assuming that scaled selection coefficients remains constant, sites with a scaled selection coefficient *γ* = 100 would take on the order of a million generations to near MSDB (see, e.g., Supplement section 2.2 in Simons et al. 2014); this roughly corresponds to 30 million years, extending back to the common ancestor of humans and Old World monkeys. With *γ* = 1, it would take on the order of 40 million generations. Disease biology and genetics has plausibly changed substantially over such time scales. We cannot rule out there being subtle footprints of asymmetry between risk-increasing and -decreasing alleles owing to fitness effects of complex diseases over some evolutionary timescales (see below). Nonetheless, such fitness effects do not explain the “smile” architecture of common variation affecting complex diseases.

What could explain the “smile” architecture is if common variants affecting disease risk were selected on because of their pleiotropic effects on myriad quantitative traits that have been subject to stabilizing selection over long evolutionary time scales (see also Koch et al. 2024). The negative relationship between the effect sizes of the variants and minor allele frequencies would arise if the effects on diseases today are positively correlated with their effects on quantitative traits that were under stabilizing selection over the molecular evolutionary timescales that shape architecture at MSDB. The approximate symmetry between risk-increasing and -decreasing alleles would be expected if mutations with small and moderate fitness effects were (approximately) equally likely to increase or decrease disease risk (because new mutations are always selected against under long-term stabilizing selection). Together these features would generate the “smile” architecture observed for many complex diseases.

This scenario seems plausible. The effects of most variants on complex traits are plausibly mediated by perturbations to the expression of genes in cellular, life history and other contexts in which expression is held close to an optimal, nonzero value by stabilizing selection. Indeed, most heritable variance in complex traits arises from common regulatory variants (Yang et al. 2010; 2011; Finucane et al. 2015) in regions with open chromatin in the cellular contexts that affect these traits (Boyle, Li, and Pritchard 2017; Sinnott-Armstrong et al. 2021; Spence et al. 2024). Moreover, there is an *a priori* expectation that genes that are expressed in a given context would have some nonzero, optimal expression level, and evidence that selection generally acts against eQTLs (Mostafavi et al. 2023). In this genic perspective, larger perturbations to expression would have greater effects on traits and would be more strongly selected against (see, e.g., Conrad et al. 2006; Glassberg et al. 2019; Zeng et al. 2023; Mostafavi et al. 2023), leading to a negative relationship between effect sizes and minor allele frequencies. The approximate symmetry between trait-increasing and -decreasing alleles would arise if weakly and moderately selected perturbations to gene expression were approximately equally likely to increase or decrease gene expression.

The kind of pleiotropic stabilizing selection that would generate the “smile” architecture has been shown to explain key features of the genetic architecture observed in GWAS of highly polygenic quantitative traits. A simple model (with few parameters) of direct and pleiotropic stabilizing selection was shown to fit the joint distribution of frequencies and effect sizes of GWAS hits for highly polygenic quantitative traits in the UKB (Simons et al. 2022). A single parameter in the model describes the coupling between the effects of the variants on the trait and on fitness. The functional form of this relationship arises from assuming that genetic variation in the trait is highly pleiotropic (Simons et al. 2018). As in the genic case described above, this functional form associates larger effects on fitness with larger effects on a trait, and mutations affecting the trait are assumed to be equally likely to increase or decrease it, giving rise to a “smile” architecture. Additionally, the high polygenicity of complex diseases and quantitative traits that GWAS revealed has been partially attributed to ‘flattening’—whereby variants whose effects on a trait exceed a small threshold value have similar expected (asymptotic) contributions to variance in the trait (O’Connor et al. 2019). Such flattening arises under direct and pleiotropic stabilizing selection (Simons et al. 2018) but does not arise under the kind of directional selection modeled here (Supplementary section S4.2).

We would expect that there to be considerable overlap between common variation affecting complex quantitative traits and complex diseases and consequently in the selection pressures that shape their genetic architecture. From a genic perspective, variation in gene expression in myriad contexts plausibly affect both. From the other end, quantitative traits like body mass index have been estimated to have mutational target sizes that exceed half of the fraction of the genome that has been estimated to be functional (Simons et al. 2022), and the high polygenicity of diseases like schizophrenia indicates similarly large target sizes (Loh et al. 2015). While the highly pleiotropic stabilizing selection model explains key observations about the genetic architecture of both complex quantitative traits and diseases, we cannot rule out there being alternative explanations for these observations. For example, a highly pleiotropic model of directional selection on traits in which the effects of variants on different traits are uncorrelated could potentially explain current observations (and could also be viewed as “apparent” stabilizing selection; Barton 1990; A. S. Kondrashov and Turelli 1992). Moreover, as we already noted, while the kind of directional selection we modeled falls short of explaining salient features of the architecture of common variation affecting complex disease susceptibility, we cannot rule out that it does have some effects on architecture.

Notably, the predictions of our model seem to be better aligned with the genetic architecture of rare variants with large effects on disease risk. These variants are too rare to be discovered in GWAS based on genotyping and imputation, and were therefore discovered using other study designs, including association and burden tests based on whole-exome sequencing (Singh et al. 2022; Palmer et al. 2022) or whole-genome sequencing in quartet families (Satterstrom et al. 2020). In these studies, rare, large effect alleles are generally found to increase disease risk as our model predicts. A caveat is that these studies have substantially greater power to identify rare risk-increasing alleles than rare risk-decreasing ones, so the observed asymmetry could also reflect an ascertainment bias. Nevertheless, the variants discovered in this way are often LoF or copy number variants that appear to be under strong purifying selection, suggesting that the asymmetry is real.

This “large-effect” mode of architecture has been found for several common, complex diseases, notably autism spectrum disorder and schizophrenia (Satterstrom et al. 2020; Singh et al. 2022). Rare, strongly deleterious alleles are much younger than the common variation identified in GWAS and are therefore more likely to reflect selection on contemporary diseases. Purifying selection on these alleles could also reflect pleiotropic selection on other traits. However, alleles that are more specific in their effect on a given complex disease are expected to contribute more to heritable variance in that disease and are therefore also more likely to be identified in mapping studies (Spence et al. 2024).

The source of selection notwithstanding, complex disease architecture including both a strongly selected “large effect” mode and a weakly selected “polygenic” mode should resemble the one that we modeled in having a fat-tailed liability distribution. Importantly, this architecture violates the normality often assumed in inference and theory (see e.g., Dempster and Lerner 1950; Falconer 1965). These departures from normality plausibly bias current estimates of the heritability of complex disease.

In particular, they would bias estimates of the proportional contributions of small and large effect variants. Total liability-scale heritability is estimated from the correlations among relatives in disease state assuming: (i) that the liability distribution is Normal in order to derive the threshold density from the prevalence (using our Eq. 30), and (ii) that variant effect sizes are sufficiently small for liability effects to equal the ratio of risk effects and threshold density (using our Eq. 8; Dempster and Lerner 1950; Falconer 1965). The contribution of small effect variants to liability-scale heritability is estimated from GWAS based on the same assumptions (Hong Lee et al. 2011). When large effect variants contribute substantially to heritability, the departure from these assumptions results in two main biases. First, the contribution of small effect variants to the liability-scale heritability is underestimated (based on either GWAS or correlations among relatives) because assuming normality when the tail is fatter leads to an overestimate of the threshold density. Second, the relative contribution of large effect variants to the total liability-scale heritability is overestimated because the linear transformation between risk and liability effects overestimates their contribution. Consequently, current estimates of the proportional contribution of large effect variants are plausibly inflated. The magnitude of these biases depends on the departures from normality, which are unknown. These biases warrant further investigation.

In summary, there is compelling evidence that heritable variation in human complex traits, including in complex disease risk, evolves under mutation-selection-drift balance (Sella and Barton 2019). Evidence from human GWAS and other study designs suggest that, at least for common genetic variation, the mode of selection in this balance is predominantly pleiotropic stabilizing selection (Simons et al. 2018; 2022; Koch et al. 2024; Spence et al. 2024). Rare genetic variation affecting complex disease susceptibility might also be shaped by directional selection of the kind we modeled here, and other modes of selection, such as balancing selection, doubtless contribute, but appear comparatively minor. More generally, as this work illustrates, by contrasting the predictions of evolutionary models with empirical findings, we can learn about the nature of selection affecting heritable variation in complex traits, and, more generally, about the evolutionary processes that shape inter-individual differences.

## Supporting information

Suppelmentary materials

## Acknowledgements

We thank Nick Barton, Magnus Nordborg, Molly Przeworski, and Himani Sachdeva for many helpful discussions and for comments on the manuscript. We also thank members of the Sella, Przeworski and Andolfatto labs at Columbia University, and the Berg, Novembre and Steinrücken labs at the University of Chicago, for feedback on the work at various stages. This work was supported by NIH F32 grant GM126787 and R35 grant GM151257 to JJB and NIH R01 grant GM115889 to GS.

## Notes

### Competing Interest Statement

The authors have declared no competing interest.

