## Supplementary material for "Mutation-selection-drift balance models of complex diseases": Suppelmentary materials

**S9 Smile plots** **38**

### S1 Notation Table

| **Symbol** | **Meaning** |
|---|---|
| $Z$ | Liability |
| $G$ | Genetic liability |
| $E$ | Environmental liability |
| $T$ | Threshold |
| $R$ | Genetic risk |
| $L$ | Mutational target size |
| $u$ | Per site mutation rate |
| $N$ | Population size |
| $\theta/2$ | Per site population-scaled mutation rate |
| $C$ | Fitness cost of disease |
| $W$ | Fitness |
| $V_A$ | Genetic variance |
| $V_E$ | Environmental variance |
| $V_P$ | Total variance |
| $h^2$ | Heritability of liability |
| $g$ | Genotype at a site |
| $a$ | Liability effect of an allele |
| $g(a)$ | Distribution of liability effect sizes |
| $\delta_R(a)$ | Risk effect of alleles with liability effect $a$ |
| $s(a)$ | Selection coefficient of alleles with liability effect $a$ |
| $x$ | Frequency of risk allele at a site |
| $F(Z)$ | Probability that an individual's liability exceeds $Z$ |
| $f(T)$ | Probability density of liability at the threshold |
| $p_+(a), p_-(a)$ | Proportion of sites with liability effect $a$ fixed for risk increasing and risk decreasing sites respectively |
| $\gamma(a)$ | Scaled selection coefficient of a site with liability effect $a$ |
| $b(a)$ | Fixation bias toward risk decreasing alleles |
| $v(a)$ | Contribution to liability variance per unit diploid mutation rate at site with liability effect $a$ |
| $\Delta_U \bar{G}$ | Mutational increase in mean liability |
| $\Delta_S \bar{G}$ | Selection response of mean liability |
| $\bar{b}$ | Mean fixation bias weighted by effect size |
| $b_T$ | Threshold bias |
| $\phi_T$ | Standardized threshold density |
| $p_s, p_l$ | Fraction of sites in two effect model that have small, large effects |

Table 1: A summary of our notation.

#### S2 Supplementary Figures

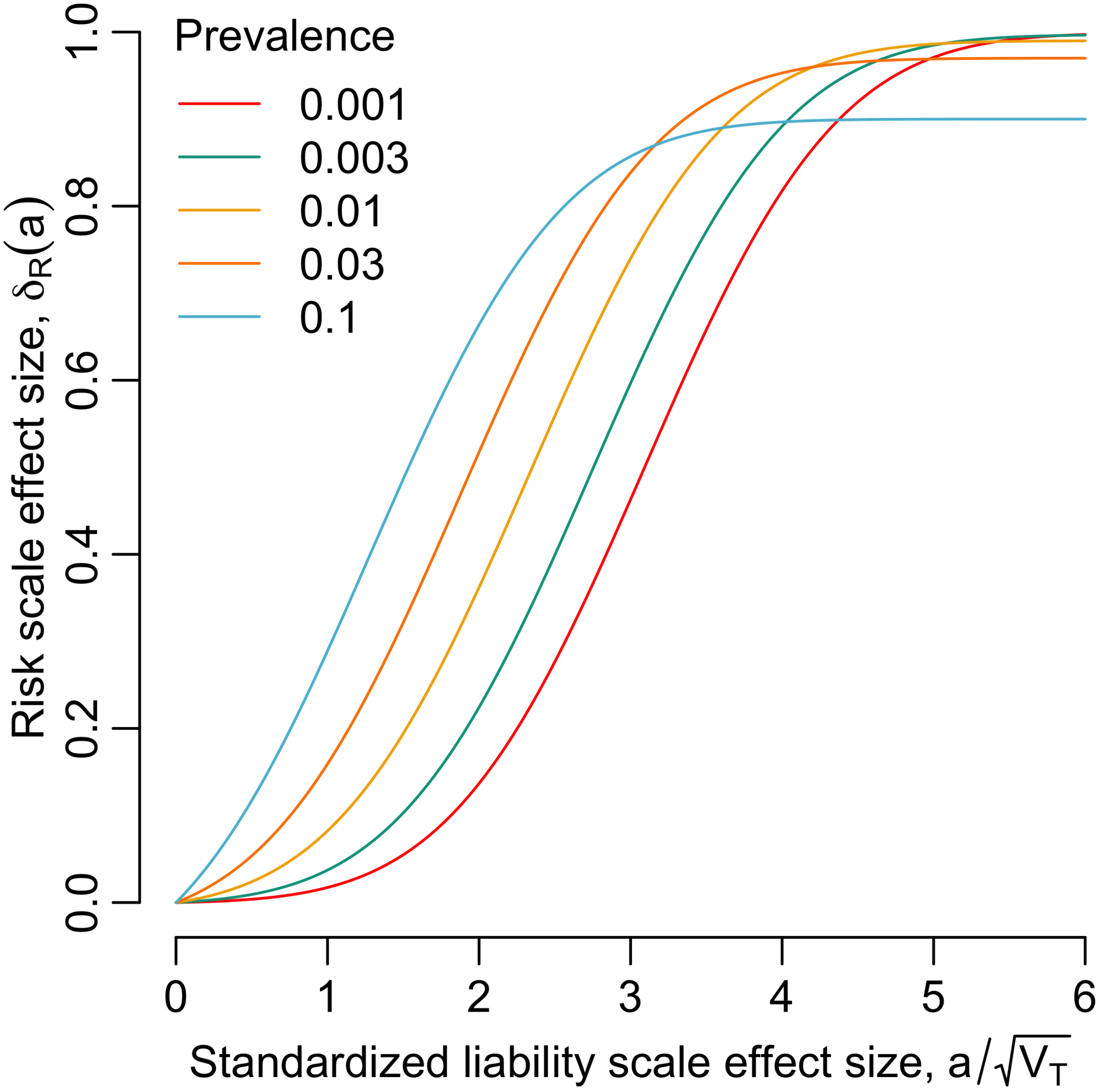

Figure S1: The risk effect as a function of the liability effect on the standardized scale. Here, we assume that the number of large effect loci is small enough that the Normal approximation applies.

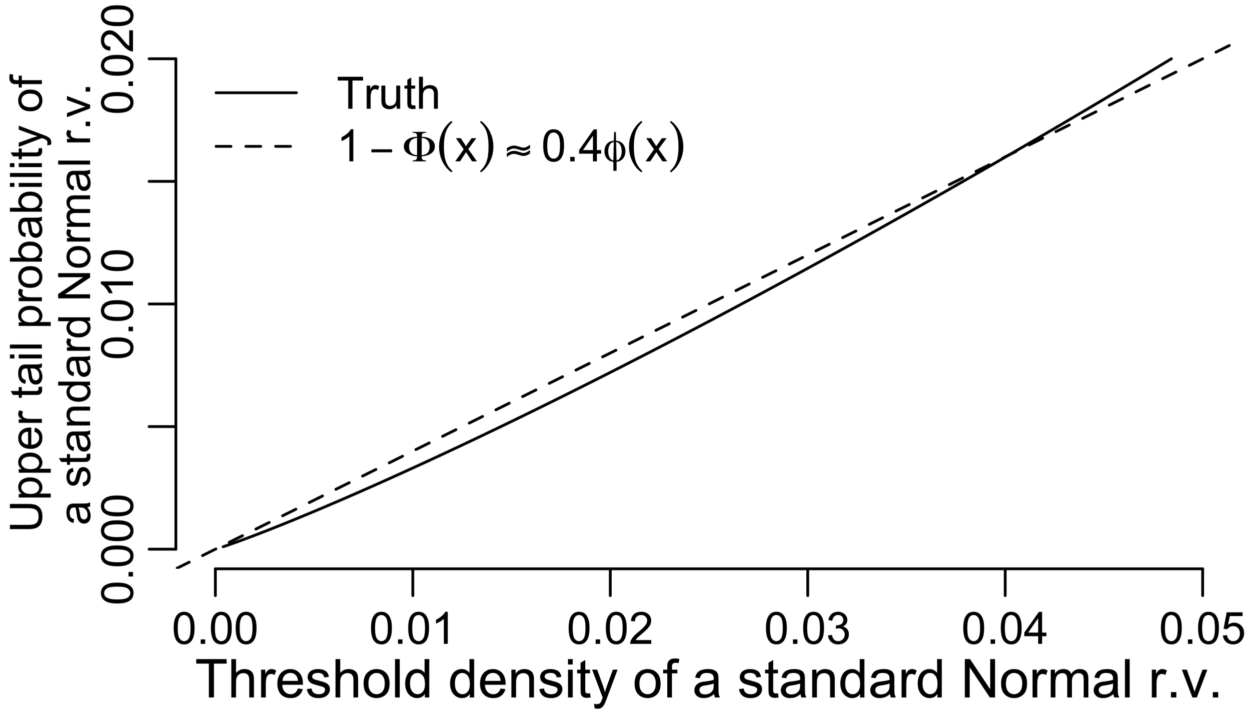

Figure S2: When the prevalence is not large, it scales approximately linearly with the threshold density.

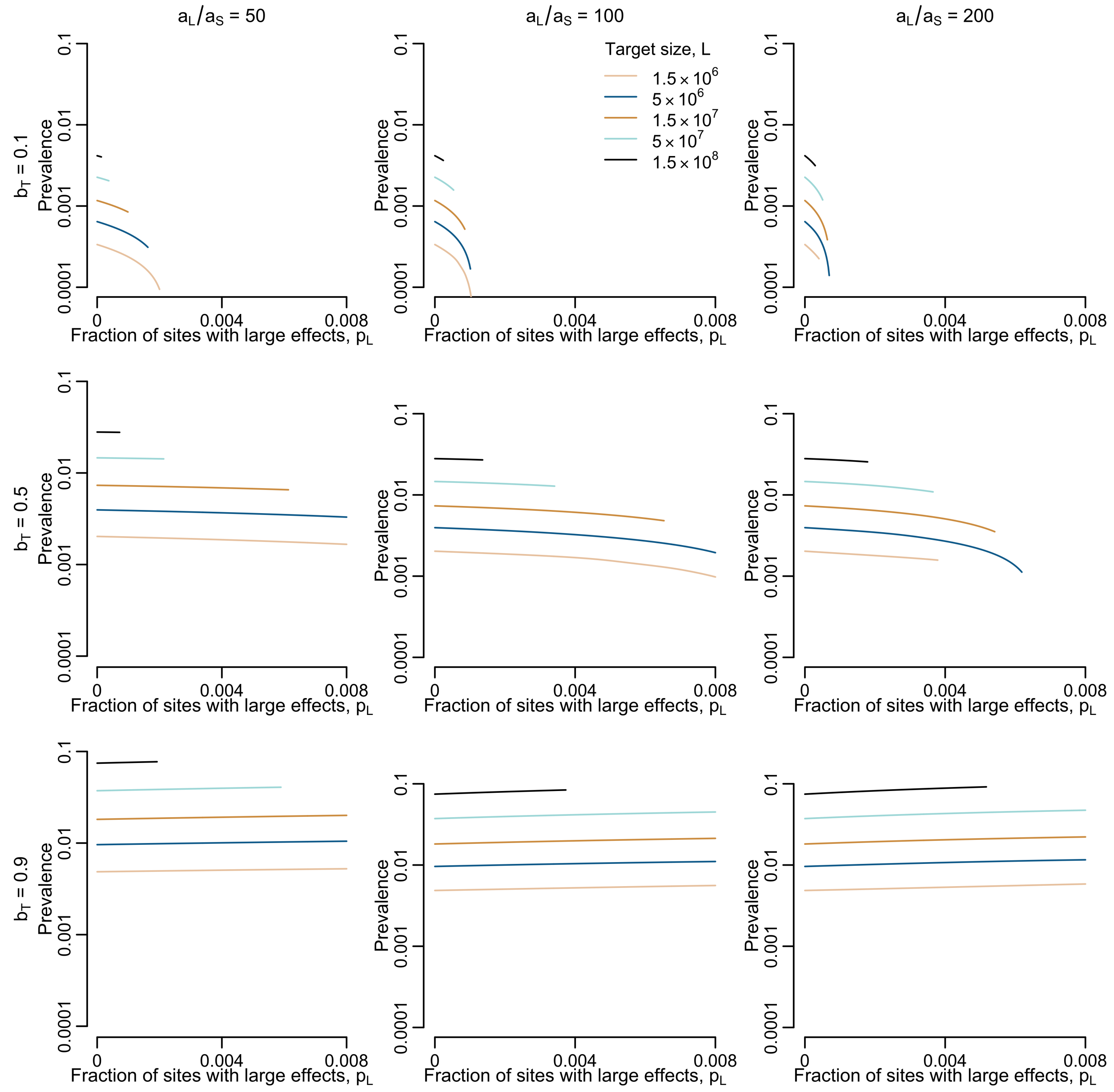

Figure S3: Each panel corresponds to a version of Figure 5B from the main text, in which we choose different values of $b_T$ and $^{a_l}/_{a_s}$ (note that we have extended the y axis so that values below 0.001 are visible). Here, the central panel corresponds to the parameters used in main text figure 5B. In general, increasing the proportion of large effect sites can cause the prevalence to go up when $b_T$ is large (though the effect is very slight), or to go down when $b_T$ is not large (in which case the effect may be more dramatic)

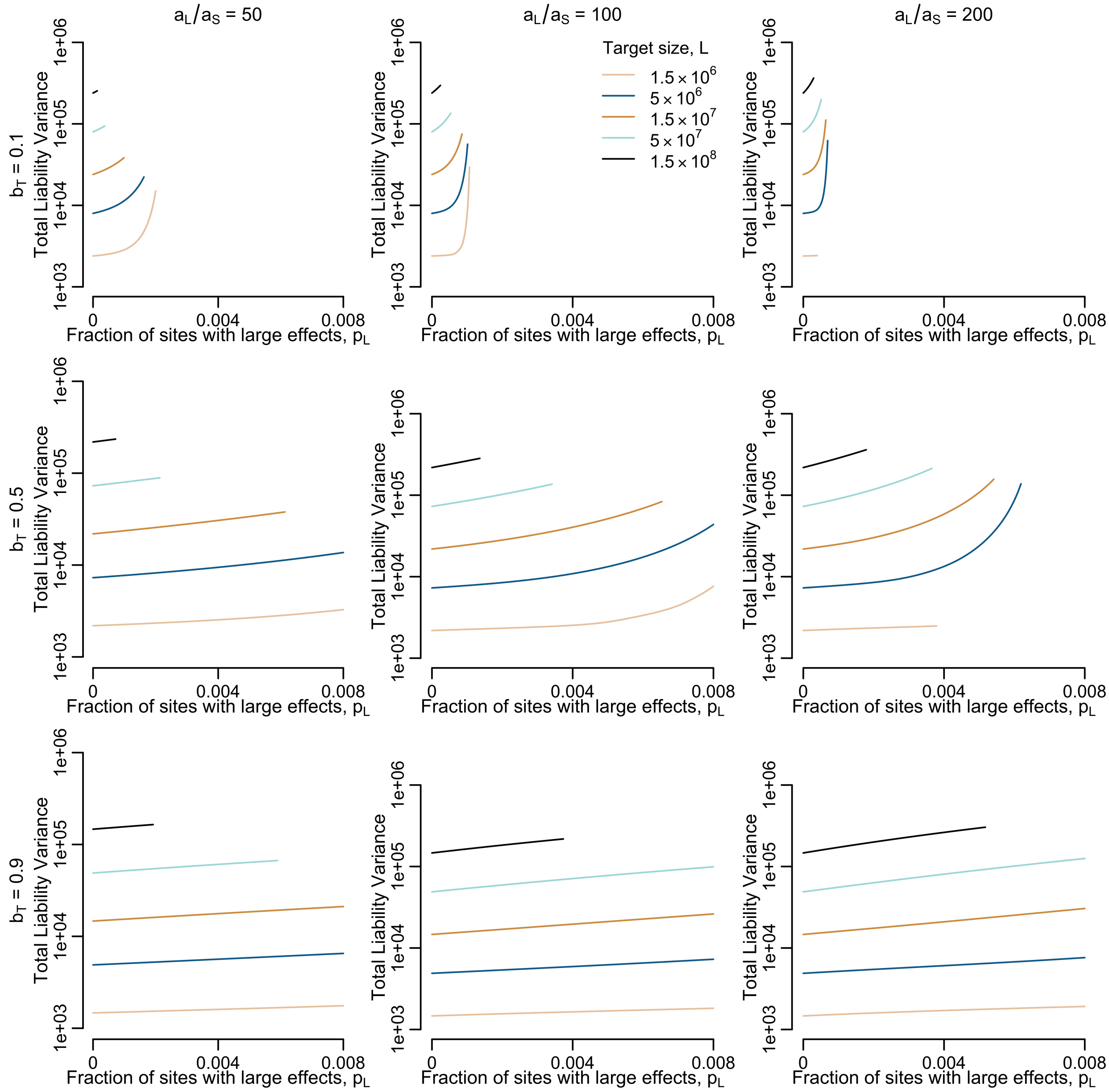

Figure S4: In each panel we plot the total variance in liability for the corresponding panel in Figure S3. As outlined in the main text, increasing the fraction of sites with large effects increases the variance, because large effect sites contribute more to variance than small effect sites. With all else held equal, an increase in the variance of liability leads to an increase in the prevalence. Comparing this figure and Figure S3, we see that when $b_T$ is large (bottom row), there is both a modest increase in the variance with increasing fraction of large effect sites, and an increase in the prevalence. In contrast, when $b_T$ is small (top row), the variance increases with an increase in the fraction of large effect sites, but the prevalence decreases.

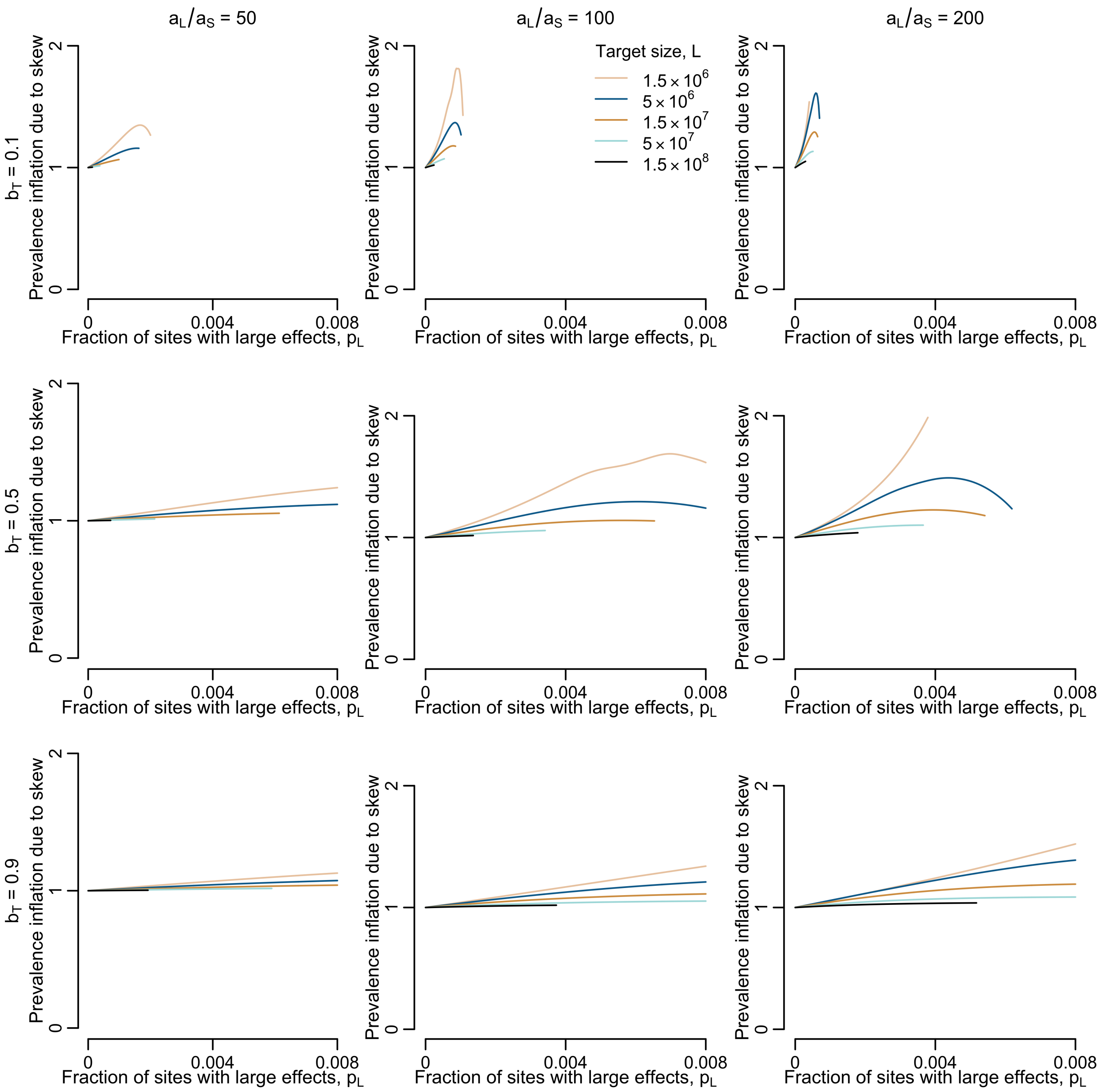

Figure S5: In each panel we plot the factor by which the prevalence is inflated due to the rightward skew of Poisson convolution distribution for the corresponding panel in Figure S3. For each predicted prevalence in Figure S3, we obtain a corresponding prevalence prediction assuming a Normal shape for the distribution, but conditional on the threshold density and variance obtained from the Poisson convolution model. We then plot the ratio of these two values. Note that this is not quite the same as fully solving the model under the Normal assumption, because we use the Poisson convolution model to obtain the large effect variance (equation (S136)). We could alternatively have substituted a Normal distribution for the Poisson convolution in equation (S134) to obtain a prevalence that "fully" assumes Normality but then inflation factors seen in this figure would also include the effect on the variance of assuming Normality when computing the strength of selection at large effect sites. This figure therefore isolates the impact of skew in the liability distribution on the prevalence, conditional on the threshold density and variance obtained from the Poisson convolution model.

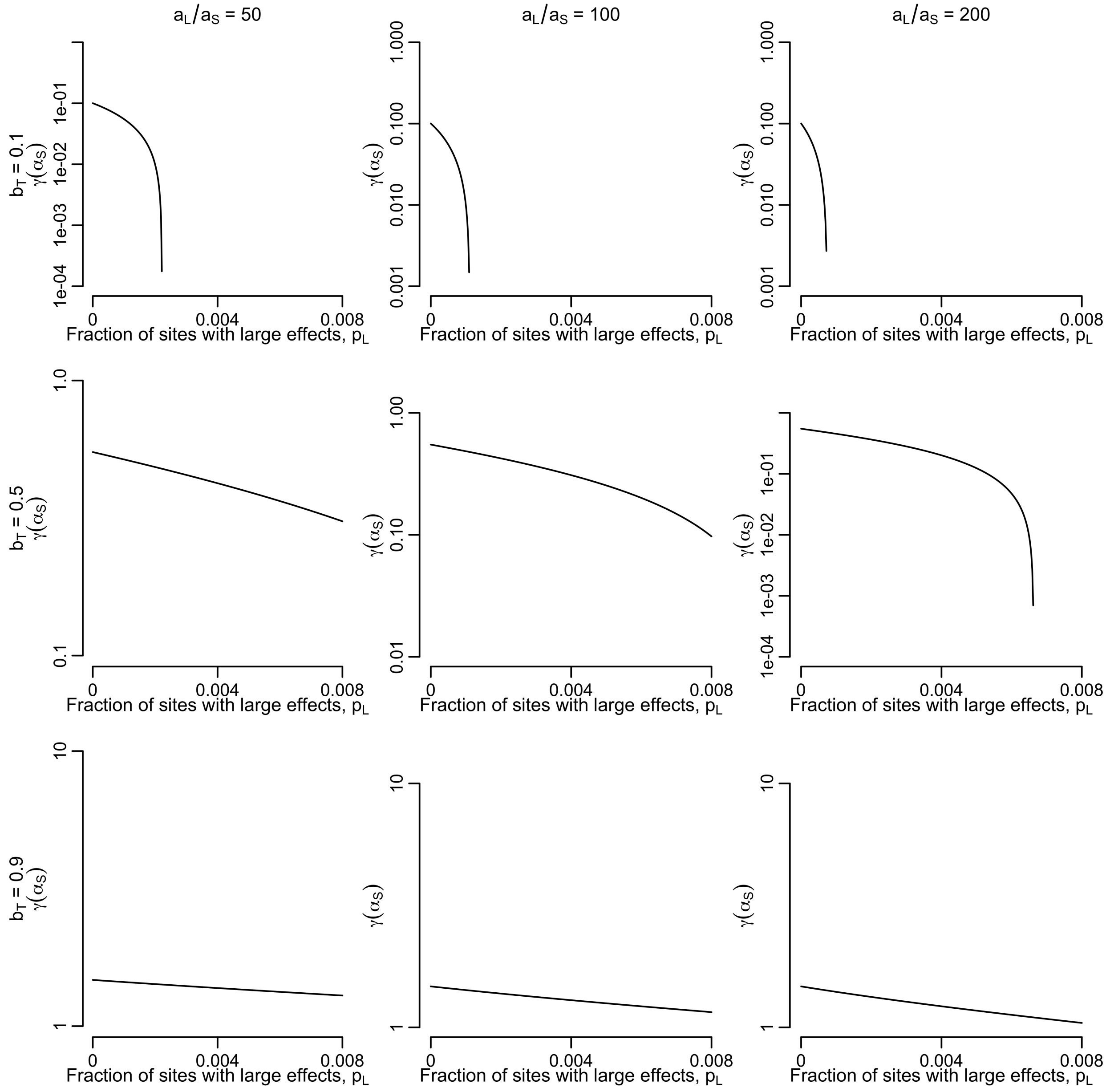

Figure S6: In each panel we plot the scaled selection coefficient for small effect sites for the corresponding panel in Figure S3. Notably, this quantity does not depend on the total number of sites, unlike the prevalence and the variance in the previous figures. In general, when we hold $b_T$ constant and increase the fraction of sites that have large effects, this leads to a decrease in the fraction of small effect sites that are fixed for the liability increasing allele, and thus a decrease in the equilibrium strength of selection acting on small effect sites.

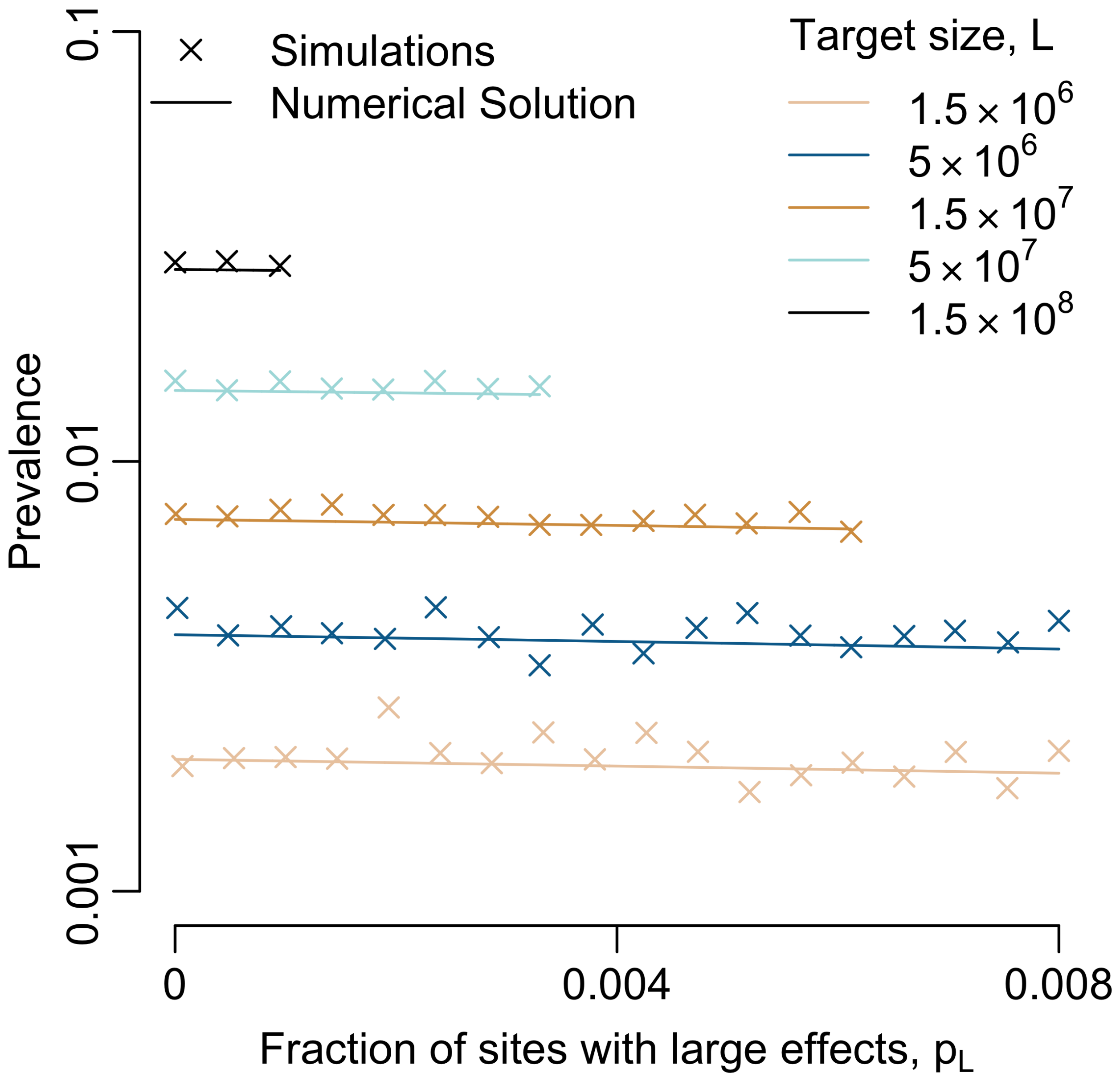

Figure S7: Simulation results match the theoretically predicted prevalence in the two effect model. Like the single effect simulations in the main text, we set $N = 1000$, $C = 0.447$, which in single effect models results in the same prevalence as $N = 20,000$, $C = 0.1$. For the other parameters, we set $b_T = 0.5$, $a_s = 1$, $h^2 = 0.5$. We also set $u = 10^{-6}$ and scale $L$ down by two orders of magnitude relative to the values in the figure legend. For the large effect sites, we set $a_L = 22.36$, which, when $p_L = 0$, yields the same value of $\delta(\alpha_L)$ with $N = 1000$, $C = 0.447$ as one obtains with $a_L = 100$ when $N = 20,000$, $C = 0.1$. However, the non-linear relationship between liability and risk for large effect sites results in a breakdown of this symmetry when $p_L > 0$, so that solutions for with $N = 1000$, $C = 0.447$, $a_L = 22.36$ and $p_L > 0$ are not the same as those for $N = 20,000$, $C = 0.1$, $a_L = 100$ and $p_L > 0$, which we plot in Figure 5B of the main text. Nevertheless, the simulations shown here serve as validation for the numerical solutions to the two effect model in a regime for which simulations are computationally feasible.

### S3 Dynamics at a site

We describe the dynamics of a site in the diffusion approximation, i.e., based on the first and second moment of changes to allele frequency in a single generation. As we note in the main text, the second moment is the standard term reflecting genetic drift. To complement the derivations in the main text, we therefore start by considering the first moment.

#### S3.1 The expected change in allele frequency

Here we derive expressions for the expected change in frequency in a single generation of an allele at frequency $x$ that increases liability by $a$. We can write the expected change at a site under viability selection as

$$\mathbb{E}\left[\Delta x\right] = -s\left(a,x\right)x\left(1-x\right) \tag{S1}$$

where

$$s\left(a,x\right) = -\frac{x\left(\overline{W}_2\left(a,x\right)-\overline{W}_1\left(a,x\right)\right)+\left(1-x\right)\left(\overline{W}_1\left(a,x\right)-\overline{W}_0\left(a,x\right)\right)}{\overline{W}}. \tag{S2}$$

The expected fitness associated with $i$ copies of the disease increasing allele at the focal site is

$$\overline{W}_i\left(a,x\right) = 1-CR_i\left(a,x\right), \tag{S3}$$

where $R_i\left(a,x\right)$ is the expected risk of individuals with $i$ copies of the risk allele. The expected fitness in the population is

$$\overline{W} = 1-C\overline{R}, \tag{S4}$$

where $\overline{R}$ is the expected disease prevalence in the population. Substituting these expressions into Eq. (S2), we find that

$$s\left(a,x\right) = \frac{C}{1-C\overline{R}}\left[x\left(R_2\left(a,x\right)-R_1\left(a,x\right)\right)+\left(1-x\right)\left(R_1\left(a,x\right)-R_0\left(a,x\right)\right)\right]. \tag{S5}$$

The different terms in this expression have useful interpretations. Notably,

$$\delta_{R,1}\left(a,x\right) = R_1\left(a,x\right)-R_0\left(a,x\right) \tag{S6}$$

$$\delta_{R,2}\left(a,x\right) = R_2\left(a,x\right)-R_1\left(a,x\right) \tag{S7}$$

are the differences in mean risk between individuals carrying 1 vs. 0 copies of the liability increasing allele, and 2 vs. 1 copies, respectively. In turn,

$$\delta_R\left(a,x\right) = x\delta_{R,2}\left(a,x\right)+\left(1-x\right)\delta_{R,1}\left(a,x\right) \tag{S8}$$

is the expected increase in individual disease risk caused by substituting a random liability decreasing at this site by a liability increasing one; for brevity, we refer to it as the additive effect on risk. Substituting these terms into Eq. (S5) gets us to Eq. 5 in the main text, i.e.,

$$s\left(a,x\right) = \frac{C}{1-C\overline{R}}\delta_R\left(a,x\right) \approx C\delta_R\left(a,x\right), \tag{S9}$$

where the latter approximation follows from our assumption that the disease is not exceedingly common.

Our expressions for the additive effect on risk (Eq. (S8)) and the selection coefficient (Eq. (S9)) suggest that the strength of selection acting on risk alleles depends on their frequency. In the sections that follow we show that when an allele has a sufficiently small effect size or is subject to strong selection the dependence on frequency is negligible. We then show that under our assumptions these two conditions cover the entire range of effect sizes, and therefore that the dependence on frequency can be ignored.

To this end we will require approximations for $\delta_R(a,x)$, $\delta_{R,i}(a,x)$ and thus for $R_i(a,x)$. We can approximate $R_i(a,x)$ by separating the liability distribution into the contributions from the focal site and from the combined genetic and environmental background. Under our assumption of high polygenicity, changes to allele frequency at the focal site do not affect the liability distribution because the genetic background evolves to compensate for them. Consequently, the probability density of the background liability $y$ is well-approximated by

$$f(y|a,x) \approx f(y+2ax), \tag{S10}$$

and the probability that background liability exceeds $y$ is well-approximated by

$$F(y|a,x) \approx F(y+2ax), \tag{S11}$$

where $f(\bullet)$ and $F(\bullet)$ denote the total stationary probability density and probability of exceeding a given value, respectively. The expected risk of an individuals with $i$ copies of the risk allele at the focal site equals the probability that the sum of contributions from the site, $i \cdot a$, and the background, $y$, exceed the threshold, $T$; namely, that $y + i \cdot a > T$ or $y > T - i \cdot a$. The expected risk of individuals with $i$ copies of the risk allele is therefore well-approximated by

$$R_i(a,x) = F(T - i \cdot a|a,x) \approx F(T - i \cdot a + 2ax). \tag{S12}$$

This way we find that

$$\delta_{R,1}(a,x) = R_1(a,x) - R_0(a,x) \approx F(T - a(1-2x)) - F(T + 2ax) \tag{S13}$$

$$\delta_{R,2}(a,x) = R_2(a,x) - R_1(a,x) \approx F(T - 2a(1-x)) - F(T - a(1-2x)). \tag{S14}$$

#### S3.2 Selection on strongly selected alleles is insensitive to their frequency

When selection on the risk increasing allele is sufficiently strong, its frequency remains tiny and selection on it nearly always occurs in individuals that are heterozygous for the allele. In this case, we can make the approximation that $x \approx 0$, so the additive effect is approximately

$$\delta_R(a,x) = x\delta_{R,2}(a,x) + (1-x)\delta_{R,1}(a,x) \tag{S15}$$

$$\approx \delta_{R,1}(a,x) \approx F(T - a(1-2x)) - F(T+2ax) \tag{S16}$$

$$\approx F(T-a) - F(T) = \delta_R(a), \tag{S17}$$

where $\delta_R(a) = F(T-a) - F(T)$ denotes this frequency independent approximation of the additive effect on risk. When this approximation applies, the selection coefficient is well-approximated by

$$s(a) = \frac{C}{1-RC}\delta_R(a) \approx C \cdot \delta_R(a). \tag{S18}$$

This approximation applies when the scaled selection coefficient $\gamma(a) = 2NC\delta_R(a) \gg 1$.

#### S3.3 Selection on small effect alleles is insensitive to their frequency

When the effect size of the risk allele is sufficiently small we can approximate its additive effect on risk by the leading term of a Taylor approximation around $a = 0$. Further noting that $\frac{\mathrm{d}F(y)}{\mathrm{d}y} = f(y)$, we find this leading term to be

$$\delta_R(a,x) \approx a \frac{\mathrm{d}\delta_R(a,x)}{\mathrm{d}a}\Big|_{a=0} = a \cdot f(T). \tag{S19}$$

This is a standard quantitative genetics approximation for the impact of truncation selection on small effect alleles (add ref to Falconer and MacKay) and the approximation we provide in Eq. (8) of the main text. Also note that when this approximation applies, our frequency independent approximation of the additive effect on risk also applies because

$$\delta_R(a) = F(T-a) - F(T) \approx a \cdot f(T) \tag{S20}$$

when $a$ is small. This validates Eq. (7) in the main text for both small effect and strongly selected alleles.

To derive the conditions in which this approximation applies, we consider the Lagrange error term for the Taylor approximation. The error can be written

$$\delta_R(a,x) - af(T) = \frac{a^2[-4(1-x)^2 x f'(T-2c(1-x)) - (1-2x)^3 f'(T-c(1-2x)) + 4(1-x)x^2 f'(T+2cx)]}{2}, \tag{S21}$$

for some $c$ satisfying $0 < c < a$. The error changes with the frequency, $x$, due to the evolution of the background, but the maximum error is attained when $x \approx 0$. We are primarily interested in the case where the effect sizes are just large enough that the first order approximation is beginning to fail. For effect sizes in this range, we can obtain an upper bound by setting $x = 0$ and $c = a$, yielding

$$\delta_R(\alpha,x) - \alpha f(T) < -\frac{\alpha^2 f'(T-a)}{2}, \tag{S22}$$

(note that $f'(x) < 0$ in the upper tail, so the right side of (S22) is positive). It follows that our small effect approximation is a good one when

$$a \ll -2\frac{f(T)}{f'(T-a)}. \tag{S23}$$

If the distribution of liability is Normal, then we can re-express (S23) in standardized units, and in terms of the standard Normal density function, as

$$a_* \ll -2\frac{\phi(T_*)}{\phi'(T_*-a_*)} \tag{S24}$$

where $a_* = a/\sqrt{V_Z}$ and $T_* = (T-\overline{Z})/\sqrt{V_Z}$ are, respectively, the effect size and distance between the threshold and the mean, in standardized units, while $\phi(\bullet)$ and $\phi'(\bullet)$ are the standard Normal pdf and its first derivative, respectively. In this general form, it is not possible to solve (S24) and isolate $a_*$ on one side of the inequality. However, we can approximate the right side of (S24) by taking another first order Taylor expansion around the point $a_* = 0$. Doing so yields

$$-2\frac{\phi(T_*)}{\phi'(T_*-a_*)} = 2\frac{e^{-a_*\left(T_*-\frac{a_*}{2}\right)}}{T_*-a_*} \approx \frac{2}{T_*} + a_*\left(\frac{2}{T_*^2-1}\right) \tag{S25}$$

Plugging this approximation into (S24) and solving for $a_*$, we find that if the distribution of liability is Normal, then the small effect approximation is valid approximately when

$$a_* \ll \frac{2T_*}{3T_*^2-2}. \tag{S26}$$

Assuming normality, $T_*$ is related to the prevalence, $\overline{R}$, via $T_* = \Phi^{-1}\left(1-\overline{R}\right)$, where $\Phi^{-1}(\bullet)$ is the inverse of the standard Normal CDF. In figure S8 we use this relationship to plot the right side of the inequality in (S26) as a function of the prevalence. Intuitively, the small effect approximation is valid over a smaller range of standardized effect sizes when the prevalence is lower.

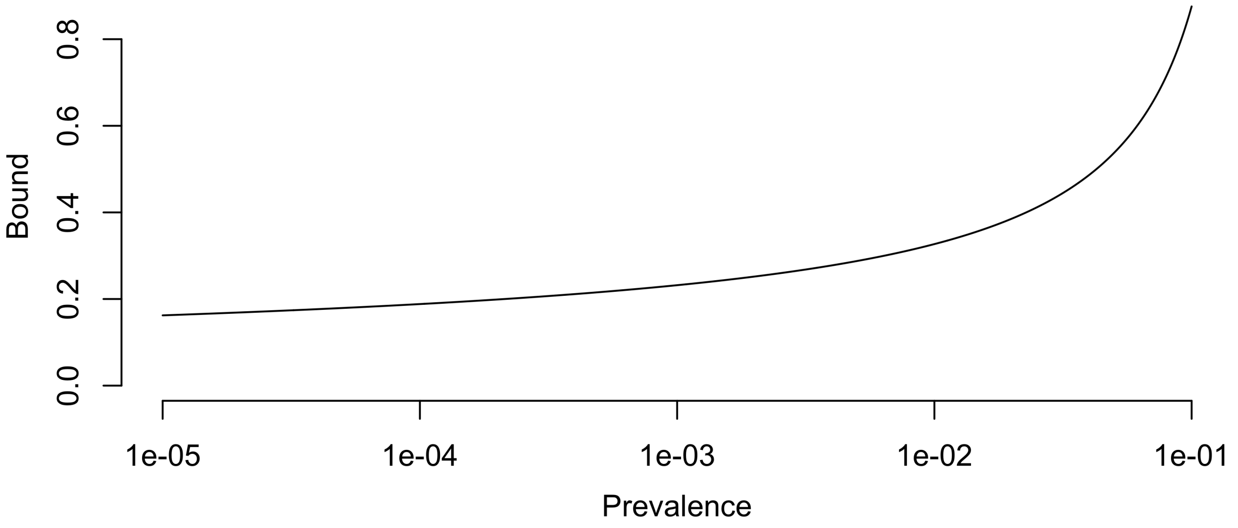

Figure S8: The upper bound on the small effect regime as a function of the prevalence, assuming that the distribution of liability is Normal.

#### S3.4 All effect sizes are small, strongly selected, or both

Here we show that under our modeling assumptions, strongly selected and small effect alleles covers the entire range of allele effects. Doing so establishes that selection on any allele in our model is insensitive to its frequency and that its selection coefficient follows one of the two approximations that we derived above.

For at least one of these approximations to apply for any allele, there must be some range of effect sizes for which both approximations apply. Sites in this range satisfy

$$\frac{1}{2Nf\left(T\right)C} \ll a \ll -2\frac{f\left(T\right)}{f'\left(T-a\right)}. \tag{S27}$$

If the liability distribution is Normal, this can restated in terms of standardized quantities, the Normal PDF, and its first derivative, as

$$\frac{1}{2N\phi\left(T_*\right)C} \ll a_* \ll -2\frac{\phi\left(T_*\right)}{\phi'\left(T_*-a_*\right)}. \tag{S28}$$

Following equation (S25), we can approximate the right term to yield

$$\frac{1}{2N\phi\left(T_*\right)C} \ll a_* \ll \frac{2}{T_*} + a_*\left(\frac{2}{T_*^2-1}\right). \tag{S29}$$

which we can rearrange to obtain

$$\frac{1}{2N\phi\left(T_*\right)C} \ll a_* \ll \frac{2T_*^2-2}{T_*^3-3T_*}. \tag{S30}$$

We can then simplify this further by removing the $a_*$ in the middle and rearranging to get

$$\frac{1}{\phi\left(T_*\right)}\frac{T_*^3-3T_*}{2T_*^2-2} \ll 2NC. \tag{S31}$$

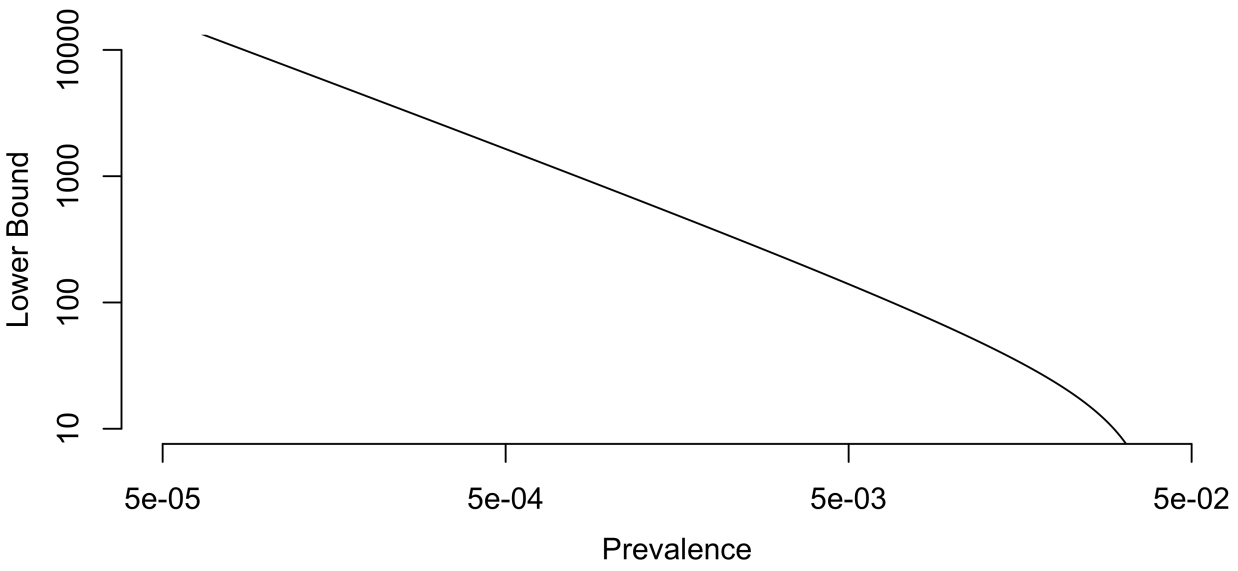

Figure S9: For a given prevalence, the strong selection and small effect regimes overlap if $2NC$ falls above the line (assuming that the shape of the liability distribution is Normal).

In figure S9 we plot the left side of (S31) as a function of the prevalence, again using the fact that $T_* = \Phi^{-1}\left(1 - \overline{R}\right)$ assuming normality. Intuitively, for a given choice of $2NC$, a relatively larger effect on liability is required for a site to be strongly selected if the prevalence is low. Thus, if a disease is quite rare and $2NC$ is not very large, then there may be sites with intermediate effects that are not covered by either of our approximations. However, for diseases and populations fitting our modeling assumptions (e.g. $\overline{R} > 1 \times 10^{-3}$, $N > 10000$, and $C > 0.1$), the condition in eq (S31) is satisfied.

If large effect sites make a substantial contribution to the variance in liability, then the distribution may not be Normal. In this case, we can write a general analog of Eq (S31) by removing the $a$ in the middle of (S27) and rearranging to obtain

$$-\frac{1}{2}\frac{f'\left(T-a\right)}{f\left(T\right)^2} \ll 2NC \tag{S32}$$

In the main text, we show that adding large effect sites has no effect on the equilibrium value of $f\left(T\right)$. It follows that Eq (S32) is a more permissive condition than Eq (S31) if $f'\left(T-a\right)$ is less negative (i.e. if the tail is flatter) in the presence of large effect sites than in their absence. This is indeed the case, so selection on individual alleles is frequency independent under our modeling assumptions in the non-Normal case as well.

#### S3.5 Relationship between the risk effect and other standard measures of risk

The risk effect of an allele, $\delta_R\left(a,x\right)$, which features throughout our derivations, is closely related to other measures used to quantify the impact of alleles on disease risk. Notably, the odds ratio of a variant with liability effect $a$ and frequency $x$ can be written as

$$OR\left(a,x\right) \equiv \frac{\text{probability of disease with the variant}}{\text{probability of disease without the variant}} = \frac{1-R}{R}\frac{R+\delta_R\left(a,x\right)}{1-\left(R+\delta_R\left(a,x\right)\right)} \tag{S33}$$

and the the risk effect can be written in terms of the odds ratio as

$$\delta_R\left(a,x\right) = -\frac{R\left(1-R\right)\left(1-OR\left(a,x\right)\right)}{1-R\left(1-OR\left(a,x\right)\right)}. \tag{S34}$$

Similarly, relative risk of the focal site can be written in terms of the risk effect as

$$RR\left(a,x\right)=\frac{R+\delta_{R}\left(a,x\right)}{R-\delta_{R}\left(a,x\right)} \tag{S35}$$

and the risk effect can be written in terms of the relative risk as

$$\delta_{R}\left(a,x\right)=R\times\frac{RR\left(a,x\right)-1}{RR\left(a,x\right)+1}. \tag{S36}$$

### S4 Genetic architecture

#### S4.1 Distribution of allele frequencies

##### S4.1.1 Taylor approximation of Wright's formula

In the Wright-Fisher diffusion for a single site under additive selection with scaled selection coefficient $\gamma > 0$ and symmetric mutation with scaled rate $\theta/2$, the probability density of the frequency of the deleterious allele is

$$\Psi\left(x \mid \theta, \gamma\right) = K_0^{-1} e^{-2\gamma x} \left(x\left(1-x\right)\right)^{\theta-1} \tag{S37}$$

where the normalization constant,

$$K_0 = \int_0^1 e^{-2\gamma x} \left(x\left(1-x\right)\right)^{\theta-1} \mathrm{d}x \tag{S38}$$

ensures that the distribution integrates to 1. We assume that the per-site population scaled mutation rate is low (i.e. $\theta/2 \ll 1$), so we can make a first order Taylor expansion around the point $\theta = 0$ to arrive at an infinite sites approximation:

$$\Psi\left(x \mid \theta, \gamma\right) \approx \psi\left(x \mid \theta, \gamma\right) = \theta \frac{d}{d\theta} \Psi\left(x \mid \theta, \gamma\right)\Big|_{\theta=0} \tag{S39}$$

$$= \frac{\theta}{1+e^{-2\gamma}} \frac{e^{-2\gamma x}}{x\left(1-x\right)}. \tag{S40}$$

Note that in this parametrization, $\gamma$ is positive, and describes the strength of selection acting against the liability increasing allele, regardless of whether it is derived or ancestral.

##### S4.1.2 An infinite sites model with two-way mutation

Although it amounts only to making approximations in a different order, it is instructive to consider an alternative derivation of equation (S40), in which we start with a more familiar infinite sites approximation, and think of the process at a given site as a combination of two processes: one that creates derived deleterious alleles, and one that creates derived beneficial alleles.

To this end, we write $q$ for the frequency of the derived allele. Following [14, 3], the derived allele at a newly arising mutation with scaled selection coefficient $\gamma$ will spend

$$\xi\left(q \mid \gamma\right) = \frac{2}{1-e^{-2\gamma}} \frac{1-e^{-2\gamma(1-q)}}{q\left(1-q\right)}. \tag{S41}$$

generations in the frequency interval $[q, q + \mathrm{d}q]$. (Note: by convention, equation (S41) is polarized so that the allele is under positive selection if $\gamma$ is positive, and negative selection if it is negative.)

The probability that the risk increasing allele is derived for any given mutation depends on the identity of the fixed allele at the site that produced the mutation. As we explain in the main text, the fixed state follows from the detailed balance equation

$$p_+\left(\gamma\right)\pi\left(\gamma\right) = p_-\left(\gamma\right)\pi\left(-\gamma\right). \tag{S42}$$

where $\pi\left(\gamma\right) = \frac{2\gamma}{1-e^{-2\gamma}}$ is the fixation rate per $2N$ mutations with scaled coefficient $\gamma$, while $p_+\left(\gamma\right)$ and $p_-\left(\gamma\right) = 1 - p_+\left(\gamma\right)$ are the fraction of sites fixed for the deleterious and beneficial alleles, respectively. Eq. (S42) is identical to Eq. (9) in the main text. There, we represent the solution in terms of the bias toward the low liability state, $b\left(\gamma\right) = p_-\left(\gamma\right) - p_+\left(\gamma\right)$. The fixed states themselves are given by

$$p_+\left(\gamma\right) = \frac{1}{1+e^{2\gamma}} \tag{S43}$$

$$p_-\left(\gamma\right) = \frac{1}{1+e^{-2\gamma}}. \tag{S44}$$

Now, averaging over the probability that the derived allele is deleterious vs beneficial, a deleterious allele with scaled coefficient $\gamma$ spends an average of

$$\begin{aligned}\tau\left(x \mid \gamma\right) &= p_+\left(\gamma\right)\xi\left(x \mid -\gamma\right) + p_-\left(\gamma\right)\xi\left(1-x \mid \gamma\right) \\ &= \frac{2}{1+e^{-2\gamma}}\frac{e^{-2\gamma x}}{x\left(1-x\right)},\end{aligned} \tag{S45}$$

generations in the $[x, x+\mathrm{d}x]$ frequency interval during its transit through the population.

Therefore, if there are $\theta/2$ new mutations per site per generation, at any given time we expect there will be a total of

$$\frac{\theta}{2}\tau\left(x \mid \gamma\right) = \frac{\theta}{1+e^{-2\gamma}}\frac{e^{-2\gamma x}}{x\left(1-x\right)} = \psi\left(x \mid \theta, \gamma\right) \tag{S46}$$

mutations in the $[x, x+\mathrm{d}x]$ frequency interval, recapitulating equation (S40).

#### S4.2 Expected heterozygosity

The expected per-site contribution to heterozygosity is

$$\mathbb{E}\left[h\left(x\right) \mid \theta, \gamma\right] = \int_0^1 h\left(x\right) \Psi\left(x \mid \theta, \gamma\right) \mathrm{d}x, \tag{S47}$$

where $h\left(x\right) = 2x(1-x)$. Under the infinite sites approximation derived in the previous section, each mutation makes an expected lifetime contribution of

$$\int_0^1 h\left(x\right) \tau\left(x, \gamma\right) \mathrm{d}x = \frac{4}{1+e^{-2\gamma}} \int_0^1 e^{-2\gamma x} \mathrm{d}x \tag{S48}$$

$$= \frac{4}{1+e^{-2\gamma}} \frac{1-e^{-2\gamma}}{2\gamma} \tag{S49}$$

$$= 2\frac{e^{2\gamma}-1}{e^{2\gamma}+1}\frac{1}{\gamma} \tag{S50}$$

$$= 2\frac{b\left(\gamma\right)}{\gamma} \tag{S51}$$

to heterozygosity during its transit through the population. The expected contribution per-site is therefore approximately

$$\mathbb{E}\left[h\left(x\right) \mid \theta \ll 1, \gamma\right] \approx \frac{\theta}{2} \int_0^1 h\left(x\right) \tau\left(x, \gamma\right) \mathrm{d}x \tag{S52}$$

$$\approx \theta \frac{b\left(\gamma\right)}{\gamma}. \tag{S53}$$

#### S4.3 Contributions to genetic variance

##### S4.3.1 Liability scale

A variant at frequency $x$ with liability effect $a$ makes contributions of $v\left(a, x\right) = a^2 h\left(x\right) = 2\alpha^2 x\left(1-x\right)$ to the genetic variance for liability. The expected per-site contribution on the liability scale is

$$\mathbb{E}[v\left(a, x\right) \mid \theta \ll 1, \gamma\left(a\right)] = a^2 \mathbb{E}[h\left(x\right) \mid \theta \ll 1, \gamma\left(a\right)] \tag{S54}$$

$$\approx a^2 \theta \frac{b\left(a\right)}{\gamma\left(a\right)} \tag{S55}$$

We can understand the expected contribution to variance by considering the three selection regimes.

For effectively neutral alleles, we can assume that both $\gamma\left(a\right) \ll 1$ and $\gamma\left(a\right) \approx 2Naf\left(T\right)C$ hold. Making a first order Taylor approximation of $b\left(a\right)$ around $a = 0$ similarly yields $b\left(a\right) \approx 2N_e \alpha f\left(T\right)C$, so that

$$\mathbb{E}[v\left(a, x\right) \mid \theta \ll 1, \gamma\left(a\right) \ll 1] \approx a^2 \theta \tag{S56}$$

for sites in the effectively neutral regime. This reflects the fact that within the effectively neutral regime, the distribution of allele frequencies is largely insensitive to differences among sites of different effect sizes in the strength of selection, so the contribution to genetic variance simply scales with the squared effect size.

In contrast, for weakly selected sites, for which $\gamma\left(a\right) \sim 1$, we can make the small effect approximation that $\gamma\left(a\right) \approx 2Naf\left(T\right)C$, but the Taylor approximation of $b\left(a\right)$ is not justified. Therefore, we have

$$\mathbb{E}[v\left(a, x\right) \mid \theta \ll 1, \gamma\left(a\right) \sim 1] \approx ab\left(a\right) \frac{2u}{f\left(T\right)C}. \tag{S57}$$

The scaling of the fixation asymmetry, $b\left(a\right)$, with the effect size $a$, is less than linear in this regime, so the variance increases more slowly than the quadratic increase it exhibits in the effectively neutral regime.

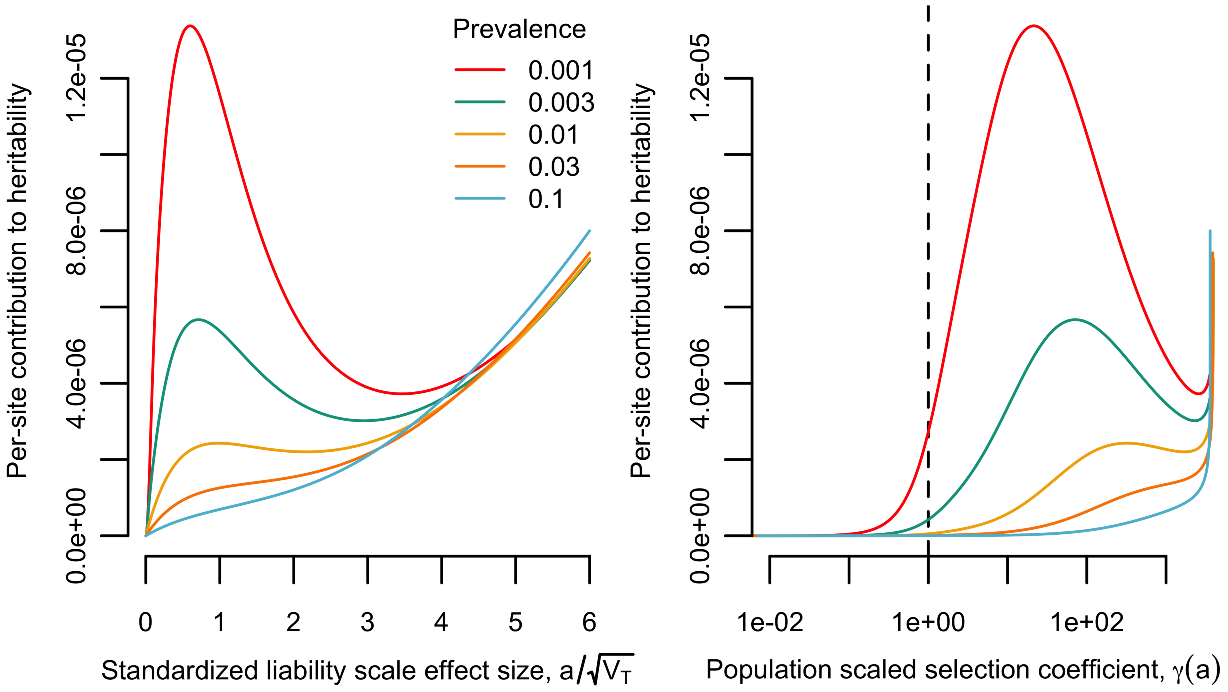

Figure S10: Left: the per site contribution to heritability as a function of the standardized liability scale effect size. Right: the same quantity, but as a function of the population scaled selection coefficient.

Finally, for strongly selected sites $\gamma(a) \gg 1$, the $\gamma(a) \approx 2N_e a f(T) C$ approximation is no longer valid either. However, here we can assume that $b(a) = 1$, so we have

$$\mathbb{E}[v(a,x) \mid \theta \ll 1, \gamma(a) \gg 1] \approx \frac{a^2}{\delta_R(a)} \frac{2u}{C}. \tag{S58}$$

Notably, the scaling of $\delta_R(a)$ with $a$ depends on details of the equilibrium, so that the exact relationship depends on details of the parameters chosen.

In the left panel of Figure S10, we plot an example in which we assume that $N = 20000$, $u = 10^{-8}$, $C = 0.1$, and that large effect alleles contribute little enough to variance that we can make the Normal approximation for the shape of the distribution. With these assumptions, the relationship between the standardized liability effect, $a/\sqrt{V_T}$, and the expected contribution to variance depends on the threshold position and the environmental variance only through their impact on the prevalence. We therefore plot the relationship between effect size and contribution to variance for several different choices of the equilibrium prevalence.

In the right panel of Figure S10, we plot the same quantity on the y-axis, but we instead plot the population scaled selection coefficient on the x axis. The dashed vertical line indicates a scaled selection coefficient of $\gamma(a) = 1$. Notably, the contribution to variance in liability does not exhibit the same asymptotic "flattening" at scaled selection coefficients above this value as is exhibited in models of stabilizing selection.

##### S4.3.2 Risk scale

Alternately, if we are interested in the contribution to additive variance on the risk scale, a variant with risk effect $\delta_R(\alpha)$ at frequency $x$ makes a contribution $v(\delta_R(\alpha), x) = 2\delta_R^2(\alpha) x(1-x)$ to the additive genetic variance for risk. Thus, in general we have

$$\mathbb{E}[v\left(\delta_R\left(a\right),x\right) \mid \theta \ll 1, \gamma\left(a\right)] = \delta_R^2\left(a\right)\mathbb{E}[h\left(x\right) \mid \theta \ll 1, \gamma\left(a\right)] \tag{S59}$$

$$\approx \delta_R^2\left(a\right)\theta\frac{b\left(a\right)}{\gamma\left(a\right)}. \tag{S60}$$

In the effectively neutral and weakly selected regimes, the risk effects scale linearly with the liability effects, so the risk scale variance exhibits the same scaling behaviors as the liability scale variance. We have

$$\mathbb{E}[v\left(\delta_R\left(a\right),x\right) \mid \gamma\left(a\right) \ll 1] = \delta_R^2\left(a\right)\mathbb{E}[h\left(x\right) \mid \theta \ll 1, \gamma\left(a\right)] \tag{S61}$$

$$\approx \delta_R^2\left(a\right)\theta \tag{S62}$$

and

$$\mathbb{E}[v\left(\delta_R\left(a\right),x\right) \mid \theta \ll 1, \gamma\left(a\right) \sim 1] \approx \delta_R\left(a\right)b\left(a\right)\frac{2u}{C} \tag{S63}$$

respectively. In the strongly selected regime, the two effect sizes diverge as the risk effects begin to increase non-linearly with the liabilty effects. Notably, however, the selection coefficients are linear in the risk effects, so the risk scale variance in the strong selection regime scales linearly with the risk effects, as

$$\mathbb{E}[v\left(\delta_R\left(a\right),x\right) \mid \theta \ll 1, \gamma\left(a\right) \gg 1] \approx \delta_R\left(a\right)\frac{2u}{C}. \tag{S64}$$

The risk variance therefore also does not exhibit flattening of the form exhibited in the stabilizing selection model. We plot the per site risk variance in figure S11, using the same parameters as we used for the liability scale variance in figure S10.

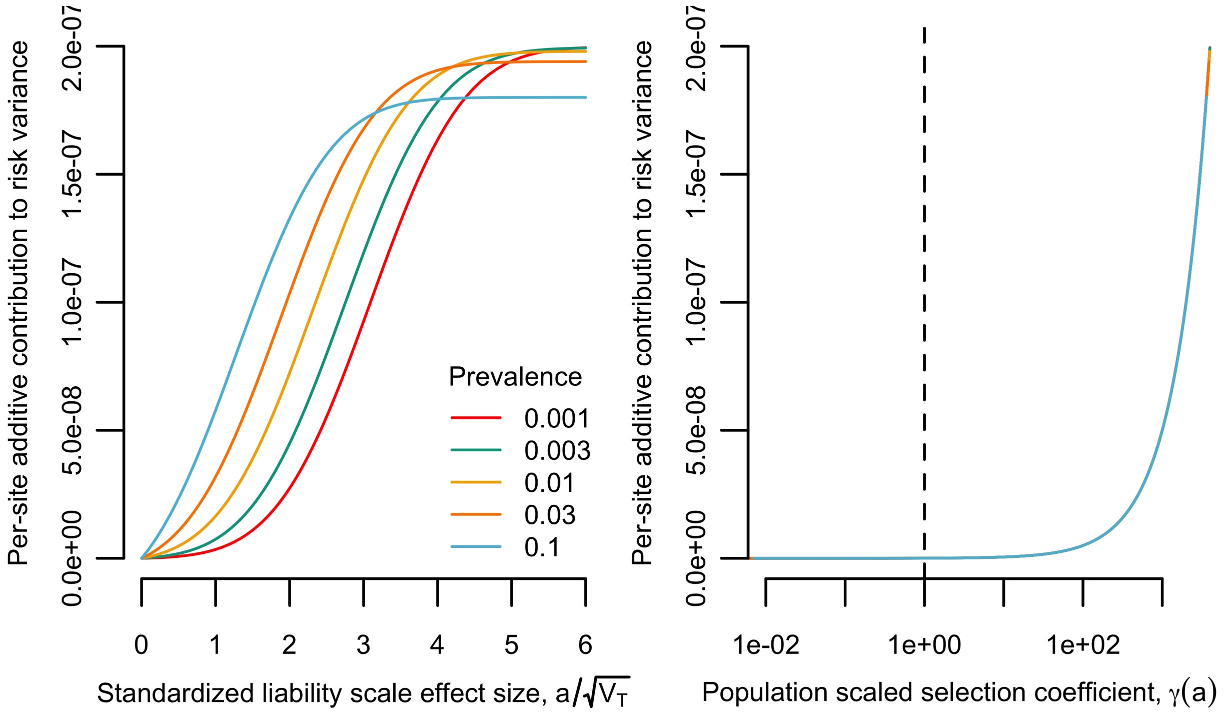

Figure S11: Left: the per site contribution to risk variance as a function of the standardized liability scale effect size. Right: the same quantity, but as a function of the population scaled selection coefficient. Note that because the risk effect and scaled selection are linearly related, and because the risk variance is also linear in the risk effect when $\gamma(a) \gg 1$, all of the lines lie on top of one another in the right panel.

#### S4.4 Signatures of asymmetry

The directional balance between mutation and selection is reflected by asymmetries in the frequency spectrum. Here we consider a few summaries of the genetic architecture which capture this asymmetry.

##### S4.4.1 Mean frequency

The mean allele frequency of the deleterious allele in the diffusion with two-way mutation and additive selection is

$$\mathbb{E}\left[x \mid \theta, \gamma\right] = \int_0^1 x \Psi\left(x \mid \theta, \gamma\right) \mathrm{d}x. \tag{S65}$$

While it is possible to express equation (S65) in terms of infinite sums, these expressions are not particularly informative. Another issue is that, by integrating from zero to one, equation (S65) includes contributions from sites at frequencies below $1/4N$ and above $1 - 1/4N$, which we think of as fixed in the analogous finite population model that the diffusion approximates.

An approximation in the infinite sites limit that we employ is given by Charlesworth and McVean [8]. Here, we provide a small generalization of their result by considering the mean frequency conditional on exceeding a threshold minor allele frequency $p^*$, as is the case for GWAS hits conditional on an effect size and study sample size [11]. Our derivation also differs slightly from theirs due to some sign flips and slight differences in the choice of integration limits, but is otherwise essentially identical.

We obtain an approximation for the mean frequency among sites with minor allele frequency greater than $p^*$ by integrating our infinite sites approximation from $p^*$ to $1 - p^*$. This gives us

$$\mathbb{E}[x|\theta \ll 1, \gamma, p^* \leq x \leq 1 - p^*] \approx \frac{\int_{p^*}^{1-p^*} x\psi\left(x \mid \theta, \gamma\right) \mathrm{d}x}{\int_{p^*}^{1-p^*} \psi\left(x \mid \theta, \gamma\right) \mathrm{d}x}. \tag{S66}$$

Equation (S66) can be re-expressed as

$$\mathbb{E}[x|\theta, \gamma, p^* \leq x \leq 1 - p^*] \approx \frac{1}{1 + \Lambda e^{2\gamma}} \tag{S67}$$

where

$$\Lambda = -\frac{\int_{2\gamma(1-p^*)}^{2\gamma p^*} \frac{e^{-t}}{t} \mathrm{d}t}{\int_{-2\gamma p^*}^{-2\gamma(1-p^*)} \frac{e^{-t}}{t} \mathrm{d}t}. \tag{S68}$$

By substituting the Taylor approximation $\frac{e^{-t}}{t} \approx 1/t - 1 + t/2$ inside of the integrals and evaluating, we have

$$\Lambda \approx -\frac{\ln\left[\frac{1-p^*}{p^*}\right] + \gamma\left(\gamma - 2\right)\left(1 - 2p^*\right)}{\ln\left[\frac{1-p^*}{p^*}\right] + \gamma\left(\gamma + 2\right)\left(1 - 2p^*\right)} \tag{S69}$$

$$\approx 1 - \frac{4\gamma\left(1 - 2p^*\right)}{\ln\left[\frac{1-p^*}{p^*}\right]}, \tag{S70}$$

where the second step involves a Taylor approximation at $\gamma = 0$. If we set $p^* = \frac{1}{4N}$, to approximate the mean frequency of all segregating variants and assume $1 - \frac{1}{4N} \approx 1 - \frac{1}{2N} \approx 1$,

$$\Lambda \approx 1 - \frac{4\gamma}{\ln\left[4N\right]} \approx e^{-\frac{4\gamma}{\ln[4N]}}, \tag{S71}$$

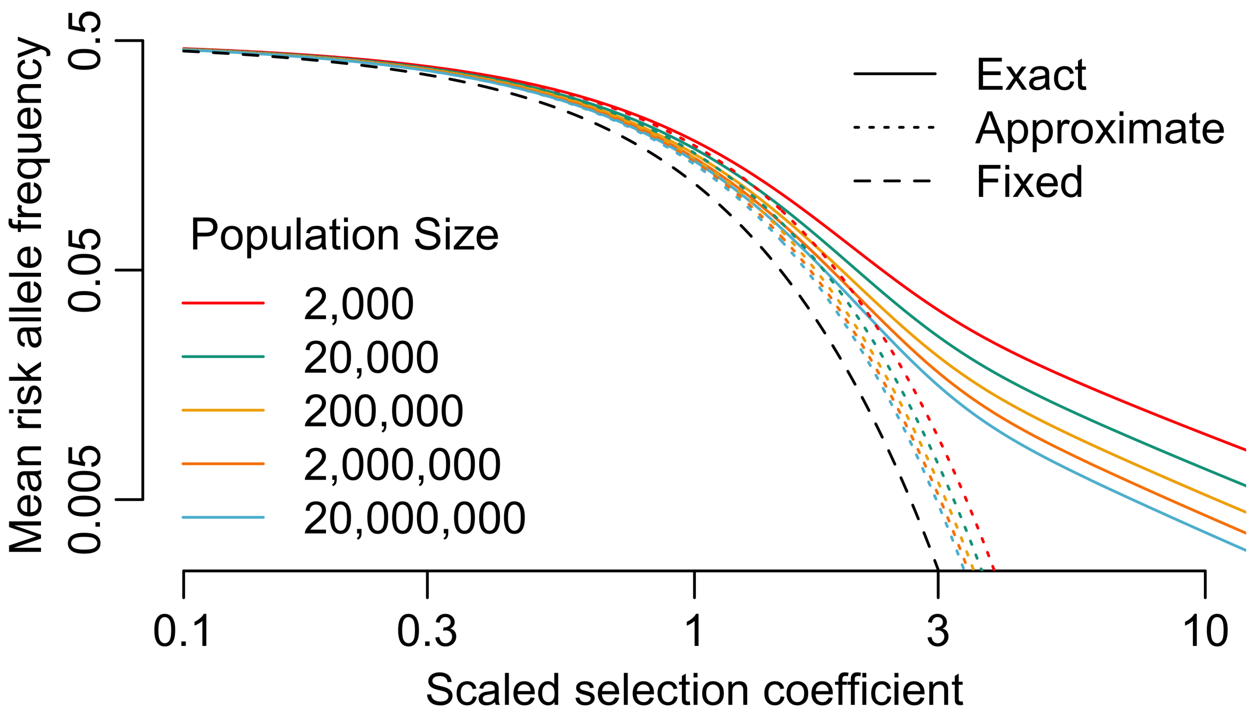

Figure S12: Mean frequency of the risk allele as a function of the scaled selection coefficient

so that

$$\mathbb{E}[x|\theta \ll 1, \gamma \lesssim 1, {}^{1}\!/_{4N} \leq x \leq 1 - {}^{1}\!/_{4N}] \approx \frac{1}{1 + e^{2\gamma(1 - {}^{2}\!/_{\ln[4N]})}}. \tag{S72}$$

which is the result of Charlesworth and McVean [8].

Thus, in large populations, the mean frequency converges to be essentially equal to the fraction of sites fixed for the beneficial allele, since

$$\lim_{N\to\infty} \mathbb{E}[x|\theta \ll 1, \gamma \lesssim 1, {}^{1}\!/_{4N} \leq x \leq 1 - {}^{1}\!/_{4N}] \approx \frac{1}{1 + e^{2\gamma}} = p_+(\gamma), \tag{S73}$$

reflecting the fact most alleles segregate at very low derived allele frequencies. We compare Eqs. (S66), (S72) and (S73) in Figure S12.

##### S4.4.2 Probability that the risk allele is derived

An additional signature of asymmetry appears in the probability that the risk allele is derived conditional on its frequency. In the infinite sites limit, this can be approximated as

$$P(\text{risk allele derived}|x, \gamma) \approx \frac{p_-(\gamma)\,\xi(x \mid \gamma)}{\tau(x \mid \gamma)} = \frac{e^{2\gamma} - e^{2\gamma x}}{e^{2\gamma} - 1} \tag{S74}$$

(Figure S13). In the neutral limit,

$$\lim_{\gamma\to 0} P(\text{risk allele derived}|x, \gamma) = 1 - x, \tag{S75}$$

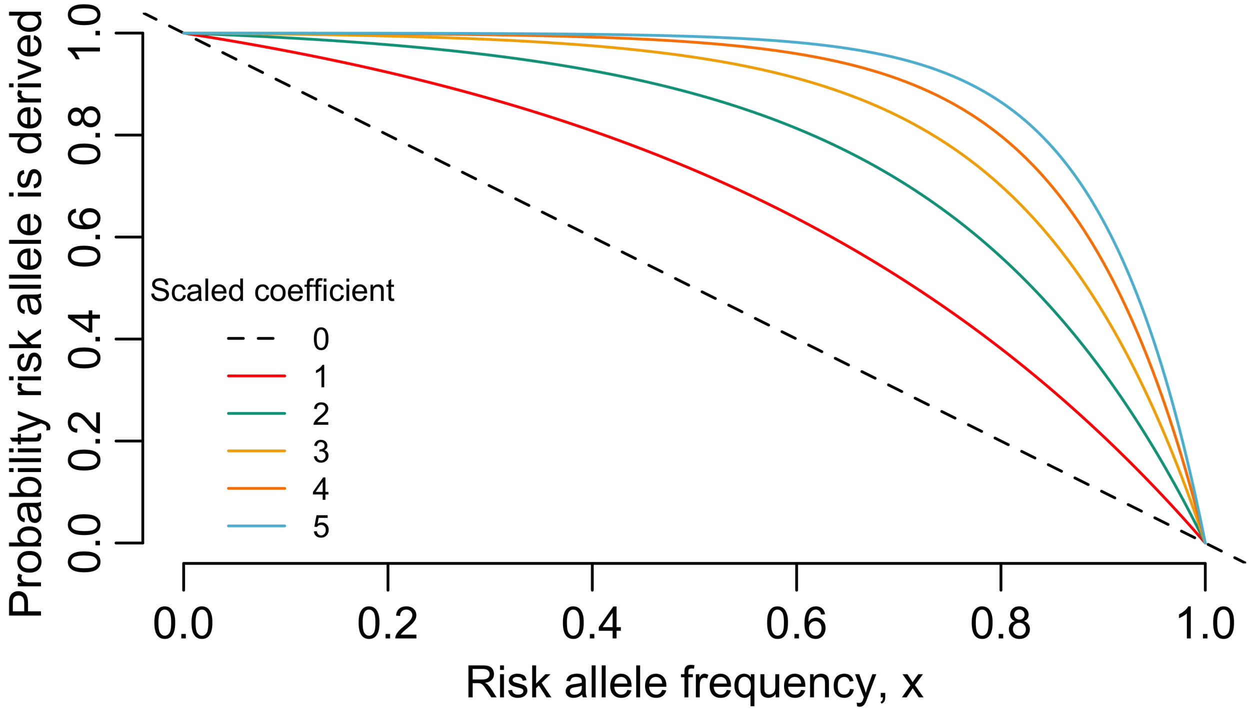

Figure S13: The probability the risk allele is derived as a function of its frequency.

whereas under strong selection,

$$P\left(\text{risk allele derived}|x, \gamma \gg 1\right) \approx 1, \tag{S76}$$

no matter the frequency, because even though risk increasing derived alleles are unlikely to reach frequency $x$ if $x$ is large, derived beneficial alleles at frequency $1-x$ are even less likely, simply because derived beneficial alleles are unlikely to exist at all. Under weak selection, $1 > P\left(\text{risk allele derived}|x, \gamma\right) > 1 - x$, with the strength of the signal scaling with $\gamma$.

### S5 Mutation and selection

#### S5.1 Exact expressions for the mutational pressure

The mutational pressure in a single generation is the difference between the expected mutational change due to liability increasing and decreasing mutations, respectively, i.e.

$$\Delta_U \overline{G} = \Delta_U^+ \overline{G} - \Delta_U^- \overline{G}. \tag{S77}$$

Liability increasing mutations occur at sites in the low liability state, and vice versa. The exact expected contribution to mutational pressure in either direction can therefore be written as a sum over the genotypes at each site

$$\Delta_U^+ \overline{G} = 2u \sum_\ell a_\ell \left(2 - g_{i\ell}\right) \tag{S78}$$

$$\Delta_U^- \overline{G} = 2u \sum_\ell a_\ell g_{i\ell}. \tag{S79}$$

where $a_\ell$ is the effect size at site $\ell$ and $g_{i\ell}$ is the genotype of individual $i$ at site $\ell$. Consolidating the sums within each effect size, we can rewrite these as

$$\Delta_U^+ \overline{G} = 2Lu \sum_a g(a) a \left(1 - \overline{x}\left(a\right)\right) \tag{S80}$$

$$\Delta_U^- \overline{G} = 2Lu \sum_a g(a) a \overline{x}\left(a\right). \tag{S81}$$

where $\overline{x}\left(a\right)$ is the mean frequency of liability increasing alleles at sites with effect size $a$ (note: both fixed and segregating sites are included in this mean). Here, $g(a)$ is interpreted as a discrete probability distribution over a finite number of effect sizes or equivalently as giving the fraction of sites with effect $a$.

Recognizing that the mean liability is $\overline{G} = 2L \sum_a g(a) a \overline{x}\left(a\right)$, we can rewrite the positive and negative components of mutational change in terms of the mean liability

$$\Delta_U^+ \overline{G} = 2Lu\overline{a} \left(1 - \frac{\overline{G}}{2L\overline{a}}\right) \tag{S82}$$

$$\Delta_U^- \overline{G} = 2Lu\overline{a} \frac{\overline{G}}{2L\overline{a}}. \tag{S83}$$

The total expected change in liability between parents and offspring due to mutation is therefore

$$\Delta_U \overline{G} = 2Lu \sum_a g(a) a \left(1 - 2\overline{x}\left(a\right)\right). \tag{S84}$$

$$= 2Lu\overline{a} \left(1 - \frac{\overline{G}}{L\overline{a}}\right). \tag{S85}$$

where these two expressions are equal to one another, and are exact under our initial, finite sites model. This justifies the statement in the main text that Eq 22 is exact when we account contributions from segregating sites.

#### S5.2 Deriving the infinite sites approximation

We now further develop the infinite sites approximation that we use in the main text.

##### S5.2.1 Site-level perspective

We first consider the sum over effect sizes. To this end, we write the mean allele frequency for sites with effect $a$ as

$$\overline{x}\,(a) = (1 - p_{seg}(a))p_{+}(a) + p_{seg}(a)\overline{x_{seg}}(a) \tag{S86}$$

where $p_{seg}(a)$ is the fraction of sites with effect $a$ that are segregating, $p_{+}(a)$ is the fraction of non-segregating sites with effect $a$ that are fixed for the liability increasing allele, and $\overline{x_{seg}}(a)$ is the mean frequency of risk increasing alleles among segregating site with effect $a$. Equation (S84) can thus be rewritten so that the mutational pressure reads

$$\Delta_U\overline{G} = 2Lu\sum_a g(a)a\left(1 - 2\left[(1 - p_{seg}(a))p_{+}(a) + p_{seg}(a)\overline{x_{seg}}(a)\right]\right) \tag{S87}$$

Now, if we take the infinite sites limit in which $L \to \infty$ and $u \to 0$ such that $\lim_{L\to\infty,u\to 0} Lu = U$, it follows that $p_{seg}(a) \to 0$, such that

$$\lim_{\substack{L\to\infty\\ u\to\infty\\ Lu=U}} \Delta_U\overline{G} = 2U\int_a g(a)a\left(1 - 2p_{+}(a)\right)\mathrm{d}a = 2U\int_a g(a)a\left(p_{-}(a) - p_{+}(a)\right)\mathrm{d}a \tag{S88}$$

$$= 2U\int_a g(a)ab(a)\mathrm{d}a \tag{S89}$$

$$= 2U\overline{a}\int_a g(a)\frac{a}{\overline{a}}b(a)\mathrm{d}a \tag{S90}$$

$$= 2U\overline{a}\int_a g(a)\frac{a}{\overline{a}}b(a)\mathrm{d}a \tag{S91}$$

$$= 2U\overline{a}\overline{b}. \tag{S92}$$

which produces equations (19) and (20) in the main text. (Note that in moving to the infinite sites limit, we have replaced the sum over a discrete number of effect sizes with an integral over a continuous distribution of effect sizes to reflect the that fact that in a model with infinite sites we can imagine the effect sizes to be continuously distributed in a way that we could not for a finite sites model.)

##### S5.2.2 Phenotypic perspective

To derive the infinite sites model from the phenotypic perspective, we rewrite the mean phenotype as the difference between the threshold position and the deviation of the mean from the threshold, i.e. $\overline{G} = T - (T - \overline{G})$, so that the mutational pressure reads

$$\Delta_U\overline{G} = 2Lu\overline{a}\left(1 - \frac{T}{L\overline{a}} + \frac{T - \overline{G}}{L\overline{a}}\right). \tag{S93}$$

The distance between the threshold and the mean is proportional to the standard deviation of total liability, i.e. $T - \overline{G} \propto \sqrt{V_T}$. The standard deviation of liability, $\sqrt{V_T}$, in turn, is proportional to $\sqrt{Lu} = \sqrt{U}$, and so it follows that $\frac{T-\overline{G}}{L\overline{a}} \to 0$ in the limit as $L \to \infty, u \to 0, Lu = U$, so the mutational pressure depends only on the relative position of the threshold, the total mutation rate, and the average effect size.

Some care must be taken with the role of the threshold in this limit. For example, if we leave the threshold fixed while taking the limit, then we have

$$\lim_{\substack{L\to\infty\\ u\to 0\\ Lu=U}} \Delta_U\overline{G} = 2U\overline{a}, \tag{S94}$$

which corresponds to the strongly selected regime, in which the fixation asymmetry is complete, and all mutations increase liability. In contrast, if we also take $T \to \infty$ at the same rate such that

$$\lim_{\substack{L\to\infty \\ T\to\infty}} (1 - \frac{T}{L\overline{a}}) = b_T \tag{S95}$$

then we have

$$\lim_{\substack{L\to\infty \\ u\to 0 \\ Lu=U \\ T\to\infty \\ (1-\frac{T}{L\overline{a}})=b_T}} \Delta_U \overline{G} = 2U\overline{a}b_T, \tag{S96}$$

which corresponds to the weakly selected regime where the fixation asymmetry is incomplete. Thus, in models with both small and large effect loci, we are from the phenotypic perspective, essentially stitching together two different infinite sites limits.

#### S5.3 Phenotypic response to selection

In the main text, we characterize the impact of selection largely terms of its impact on the long term behavior of individual sites. In this section, for completeness, we describe the phenotypic response to selection in terms of classic quantitative genetic expressions. We first give a general characterization in terms of selection acting directly on genetic risk, before showing how this can be approximated in terms of a model in which selection acts directly on liability when effects are small. We give these expressions in terms of the finite sites model.

##### S5.3.1 General expression in terms of selection acting on genetic risk

The expected fitness of individual $j$ given their genetic liability is

$$\mathbb{E}\left[W_j \mid G_j\right] = 1 - CR_j. \tag{S97}$$

By Robertson's secondary theorem [10, 9], the expected change in liability due to selection is

$$\Delta_S \overline{G} = \frac{1}{\overline{W}} Cov\left(W, G\right).$$

The mean fitness is $\overline{W} = 1 - C\overline{R}$, where $\overline{R} = F\left(T\right)$ is the mean risk, i.e. the prevalence. The expected change can therefore be written in terms of risk and cost as

$$\Delta_S \overline{G} = -\frac{C}{1 - C\overline{R}} Cov\left(R, G\right). \tag{S98}$$

Equation (S98) can be seen as a form of the multivariate breeder's equation with liability and risk being viewed as correlated traits: selection acts on risk with selection gradient $-\frac{C}{1-C\overline{R}}$. There is no "direct" selection on liability, but it exhibits a correlated response given the selection on risk, and the magnitude of this response depends on the genetic covariance between liability and risk, $Cov\left(R, G\right)$.

**Quantitative genetic decomposition of risk** The covariance term in equation (S98) depends on the details of the relationship between liability and risk. Notably, conditional on the equilibrium liability distribution, the relationship between these two quantities is deterministic, because the risk effects are simply non-linear transforms of the liability effects. To better understand, it is helpful to rewrite an individual's risk in terms of the standard quantitative genetic decomposition in which the phenotype is decomposed as a sum of orthogonal components. To this end, we write individual $i$'s genetic risk as

$$\begin{aligned} R_i = \overline{R} + \sum_{\ell} \delta_R\left(a_\ell, x_\ell\right)\left(g_{i\ell} - 2x_\ell\right) + \sum_{\ell} \zeta_R\left(a_\ell, x_\ell\right)\left(d_{i\ell} - 2h\left(x_\ell\right)\right) \\ + \sum_{\ell \neq \ell'} \eta_R\left(a_\ell, a_{\ell'}, x_\ell, x_{\ell'}\right)\left(g_{i\ell} - 2x_\ell\right)\left(g_{i\ell'} - 2x_\ell\right) + \cdots \end{aligned} \tag{S99}$$

where $\delta_R(a_\ell, x_\ell)$ and $\zeta_R(a_\ell, x_\ell)$ are the additive and dominance effects on risk for a variant with effect liability effect $a_\ell$ and frequency $x_\ell$, and $\eta_R(a_\ell, a_{\ell'}, x_\ell, x_{\ell'})$ is the additive-by-additive interaction effect between two such variants. The $\cdots$ represent additional terms, including additional forms of pairwise interaction (i.e. "additive-by-dominance" and "dominance-by-dominance" interactions), as well as interactions of higher order (e.g. three-way, four-way, etc.). Notably, because the risk, $R_j = 1 - \Phi(T \mid G_j, V_E)$, is not a polynomial function of $G_j$, all higher order terms technically make non-zero contributions to the risk. Many of these contributions will be individually small, but in aggregate they dominate the additive contribution, such that most of the genetic variance in risk is non-additive. However, liability itself is a purely additive quantity, so all of the covariances between the genetic component of liability and the non-additive components of risk are zero. As a result, $Cov(R, G)$ depends only on the additive component of variation in liability/risk, i.e.

$$\begin{aligned}
Cov(R,G) &= Cov\left(\sum_\ell \delta_R(\alpha_\ell, x_\ell)\, g_{i\ell} + \sum_\ell \zeta_R(\alpha_\ell, x_\ell)\, d_{i\ell} + \sum_{\ell \neq \ell'} \eta_R(\alpha_\ell, \alpha_{\ell'}, x_\ell, x_{\ell'})\, g_{i\ell} g_{i\ell'} + \cdots, \sum_\ell \alpha_\ell g_\ell\right) \\
&= \sum_\ell \alpha_\ell \delta_R(\alpha_\ell, x_\ell)\, Cov(g_\ell, g_\ell) + \sum_\ell \alpha_\ell \zeta_R(\alpha_\ell, x_\ell) \underbrace{Cov(d_\ell, g_\ell)}_{0} \\
&\qquad + \sum_\ell \alpha_\ell \eta_R(\alpha_\ell, \alpha_{\ell'}, x_\ell) \underbrace{Cov(g_{i\ell} g_{i\ell'}, g_\ell)}_{0} + \underbrace{\cdots}_{0} \\
&= 2 \sum_\ell \alpha_\ell \delta_R(\alpha_\ell, x_\ell)\, x_\ell (1 - x_\ell).
\end{aligned} \tag{S100}$$

The response to selection is therefore

$$\Delta_S \overline{G} = -2 \frac{C}{1 - C\overline{R}} \sum_\ell \alpha_\ell \delta_R(\alpha_\ell, x_\ell)\, x_\ell (1 - x_\ell). \tag{S101}$$

#### S5.4 Univariate breeder's equation/small effect approximation

The derivation above is general, and makes no strong assumptions beyond those outlined in the Model section in the main text. If assume that the distribution is Normal, then we can obtain a slightly different expression, which can be understood in terms of the univariate breeder's equation, with selection acting directly on liability.

The univariate breeder's equation says that the response to selection is equal to the heritability times the selection differential, $\Delta_S \overline{G} = h^2 S$. The selection differential is the difference between the mean phenotype of the parents before selection ($\overline{Z}$) and the mean after selection ($\overline{Z}'$)

$$S = \overline{Z}' - \overline{Z}. \tag{S102}$$

Under our model, the mean after selection is

$$\overline{Z}' = \frac{\left(1 - \overline{R}\right) \overline{Z}_{Z<T} + \overline{R}(1 - C)\, \overline{Z}_{Z>T}}{\left(1 - \overline{R}\right) + \overline{R}(1 - C)} \tag{S103}$$

$$= \frac{\left(1 - \overline{R}\right) \overline{Z}_{Z<T} + \overline{R}(1 - C)\, \overline{Z}_{Z>T}}{1 - \overline{R}C} \tag{S104}$$

where $\overline{Z}_{Z<T}$ is the mean liability conditional on not having the disease, and $\overline{Z}_{Z>T}$ conditional on having it. Assuming that (total) liability is Normally distributed, these quantities are

$$\overline{Z}_{Z<T} = \overline{Z} - \frac{f(T)\, V_Z}{1 - \overline{R}} \tag{S105}$$

and

$$\overline{Z}_{Z>T} = \overline{Z} + \frac{f(T) V_Z}{\overline{R}}, \tag{S106}$$

where $f(T)$ is the density of liability at the truncation point, and $V_Z$ is the total variance of liability. The selection differential is therefore

$$\begin{aligned} S &= \overline{Z}' - \overline{Z} && \text{(S107)} \\ &= \frac{\left(1-\overline{R}\right)\overline{Z}_{Z<T} + \overline{R}\left(1-C\right)\overline{Z}_{Z>T}}{1-\overline{R}C} - \overline{Z} && \text{(S108)} \\ &= \frac{\left(1-\overline{R}\right)\overline{Z} - f(T) V_Z + \overline{R}\left(1-C\right)\overline{Z} + \left(1-C\right) f(T) V_Z - Z\left(1-\overline{R}C\right)}{1-\overline{R}C} && \text{(S109)} \\ &= -\frac{f(T) C V_Z}{1-\overline{R}C} && \text{(S110)} \end{aligned}$$

and the response to selection is

$$\Delta_S \overline{G} = -h^2 \frac{f(T) C V_Z}{1-\overline{R}C}. \tag{S111}$$

We can also rewrite this in terms of the selection gradient (i.e. $\Delta_S \overline{G} = V_G \beta$, where $V_G$ is the genetic variance in liability, and $\beta = \frac{\mathrm{d} \ln \overline{W}}{\mathrm{d}\overline{Z}} = \frac{1}{\overline{W}} \frac{\mathrm{d}\overline{W}}{\mathrm{d}\overline{Z}}$ is the selection gradient). Substituting $h^2 = \frac{V_G}{V_Z}$ and canceling the $V_Z$s we have

$$\Delta_S \overline{G} = -V_G \frac{f(T) C}{1-C\overline{R}} \approx -V_G f(T) C \tag{S112}$$

where the genetic variance for liability is

$$V_G = 2 \sum_{\ell} a_{\ell}^2 x_{\ell} \left(1 - x_{\ell}\right). \tag{S113}$$

Comparing equation (S112) to equation (S98), we see that assuming Normality of the distribution of liability and applying the univariate breeder's equation is equivalent to approximating the risk effect as $\delta_R(\alpha) \approx \alpha f(T)$, i.e. the small effect approximation which we developed in Section S3.

### S6 Effect size variation

When there is a distribution of effect sizes, the equilibrium satisfies

$$b_T = \int_a b(a) \frac{a}{\overline{a}} \mathrm{d}a \tag{S114}$$

In this section, our goal is to understand how variation in the distribution of effect sizes among small effect sites will impact the equilibrium. To this end, we compare two models, one in which there is only a single effect size, and one where the effects follow some arbitrary distribution $g(a)$, assuming that the sites in the upper tail of the distribution still fall within the small effect regime.

We use a subscript $g$ to denote quantities associated with the model with a distribution of effects, and a subscript $se$ to denote those associated with the single effect model. For example, $\alpha_{se}$ is the effect size in the single effect model, while $\overline{a_g} = \int_a g(a)\, a da$ is the mean effect given that effects come from the distribution $g$ (in general, by writing the subscript $g$ under the bar, we mean that relevant average includes an average over the distribution of effects).

We assume all other parameters are the same between the two models, and that the units of liability in the model with a distribution are chosen so that the mean effect size in that model is equal to the effect in the single effect model, i.e. $a_{se} = \overline{a_g}$. Given this assumption, the mutational pressure is the same between the two models, so it follows that the selection response must be as well, i.e.

$$\Delta_S \overline{G_g} = \Delta_S \overline{G}_{se}. \tag{S115}$$

The form of the selection response (Eq. (S112)) and the linear relationship between the scaled selection coefficient and threshold density among small effect sites (i.e. $\gamma(a) = 2Naf(T)C$) implies that

$$\frac{\overline{\gamma(a)_g}}{\gamma(a)_{se}} = \frac{f_g(T)}{f_{se}(T)} = \frac{V_{G,se}}{V_{G,g}}. \tag{S116}$$

Thus, as we would expect, if differences in the distribution of effects lead to an increase (decrease) in the mean scaled selection coefficient (and consequently, the threshold density), they also lead to a decrease (increase) in the genetic variance.

To gain some further insight, it is useful to first rewrite the genetic variance in a different way. First, recall from the main text that $v(a)$ is the contribution to variance per unit mutation rate at sites with effect $a$. In the model with a distribution of effects, the genetic variance can be written as

$$V_{G,g} = 2NLu\overline{v(a)_g} \tag{S117}$$

$$= 4NLu\overline{a_g^2} r(b_T)_g \tag{S118}$$

$$= 4NLu\overline{a_g}^2 \left(1 + CV_g^2\right) r(b_T)_g \tag{S119}$$

where $0 < r(b_T)_g = \frac{\overline{v(a)_g}}{2\overline{a_g^2}} < 1$ describes the extent to which the variance is reduced by selection relative to what it would otherwise be under neutrality given the value of $b_T$ and the distribution of effects, $g$. When $b_T \approx 0$, selection is ineffective, so $r(b_T)_g \approx 1$. The value of $r(b_T)_g$ then decreases as $b_T$ approaches 1. To arrive at equation (S119), we use the fact that $\overline{a_g^2} = \overline{a_g}^2 \left(1 + CV_g^2\right)$, where $CV_g^2 = \frac{Var_g(a)}{\overline{a_g}^2}$ is the squared coefficient of variation of the effect size distribution.

For the single effect model, the genetic variance can be written analogously as

$$V_{G,se} = 2NUv(a_{se}) \tag{S120}$$

$$= 4NUa_{se}^2 r(b_T)_{se}. \tag{S121}$$

where $0 < r\left(b_T\right)_{se} = \frac{v(a)_{se}}{2a_{se}^2} < 1$ similarly describes the magnitude of the variance reduction due to selection in the single effect model. We assumed above that the units are chosen so that $a_{se} = \overline{a_g}$. It follows that $a_{se}^2 = \overline{a}_g^2$, so in comparing the two models, we find that

$$\frac{\overline{\gamma_g\left(a\right)}}{\gamma_{se}\left(a\right)} = \frac{f_g\left(T\right)}{f_s\left(T\right)} = \frac{V_{G,se}}{V_{G,g}} = \frac{v\left(b_T,g\right)}{\left(1+CV_a^2\right)} \tag{S122}$$

where $v\left(b_T,g\right) = \frac{r_{se}\left(b_T\right)}{r_g\left(b_T\right)}$ is the ratio of the variance reduction due to selection in the single effect model to that in the distribution model.

To understand the effect on the prevalence, we recall from the main text that the standardized density, $\varphi_T = f(T)\sqrt{V_G}$, and that the prevalence scales roughly linearly with the threshold density in the range that we are interested in. Thus variation in the effect distribution affects the prevalence as

$$\frac{\overline{R_g}}{R_{se}} \approx \frac{\varphi_{T,g}}{\varphi_{T,se}} = \frac{f_g\left(T\right)\sqrt{V_{G,g}}}{f_s\left(T\right)\sqrt{V_{G,se}}} = \sqrt{\frac{V_{G,se}}{V_{G,g}}} = \sqrt{\frac{v\left(b_T,g\right)}{\left(1+CV_a^2\right)}} \tag{S123}$$

##### S6.1 Numerical solutions assuming a gamma distribution

To better understand the implications of equations (S122) and (S123), we solve the model numerically, assuming a gamma distribution for the distribution of effect sites $g$. Although we present the solution in the main text in terms of solving for $f(T)$, it turns out to be convenient here to solve the model in a different way. Specifically, we assume that effect sizes are measured in the same units as the scaled selection coefficients, i.e. $a = \gamma(a) = \gamma$ This choice of units fixes the threshold density at $f(T) = 1/2NC$ and converts the problem of solving for the threshold density into one of solving for the mean scaled selection coefficient. We then assume that $\gamma \sim \Gamma(\alpha,\beta)$ with shape and rate parameters $\alpha$ and $\beta$, and pdf $f_\Gamma(\gamma \mid \alpha,\beta)$. The shape and rate parameters of a gamma distribution are related to its mean $(\overline{\gamma})$ and coefficient of variation $(CV_\gamma)$ via

$$\alpha = CV_\gamma^{-2} \tag{S124}$$
$$\beta = \overline{\gamma}^{-1}CV_\gamma^{-2}. \tag{S125}$$

Thus, writing $b(\gamma) = \frac{e^{2\gamma}-1}{e^{2\gamma}+1}$ for the fixation bias of sites with scaled coefficient $\gamma$, we solve

$$0 = \overline{\gamma}b_T - \int_0^\infty f_\Gamma(\gamma \mid CV_\gamma^{-2}, \overline{\gamma}^{-1}CV_\gamma^{-2})\gamma b(\gamma)\mathrm{d}\gamma \tag{S126}$$

for the mean $\overline{\gamma}$, given a fixed choice of $b_T$ and $CV_\gamma^2$. We plot the $\overline{\gamma}$ obtained from these solutions in the left panel of figure S14. In the right panel, we plot equation (S122). In Figure S15, we plot equation (S123).

Visualizing these solutions helps provide intuition about equations (S122) and (S123). When $b_T \approx 0$, all sites belong to the effectively neutral regime, so $v\left(b_T,g\right) \approx 1$. The impact of variation in the effect size distribution is therefore to increase the genetic variance by a factor of $\left(1+CV_\gamma^2\right)$, and consequently to decrease the mean scaled selection coefficient by a factor of $\left(1+CV_\gamma^2\right)^{-1}$. However, if we consider larger values of $b_T$, the role of selection becomes more important. In this case, $v\left(b_T,g\right) > 1$, because sites with larger effect sizes have larger reductions in variance. In general, we see that $v\left(b_T,g\right)$ increases monotonically with $b_T$, until it eventually exceeds $\left(1+CV_\gamma^2\right)$, so that as $b_T$ approaches 1, the total effect of variation in the effect distribution is to decrease the genetic variance, and therefore to increase the mean scaled selection coefficient. Curiously, when the coefficient of variation of $g$ is close to zero, this crossover happens when $\overline{\gamma}b_T \approx 1$ (i.e. at approximately $\overline{\gamma} \approx 1.12$ and $b_T = 0.83$). For distributions with a larger variance, the cross over happens at higher values of $b_T$.

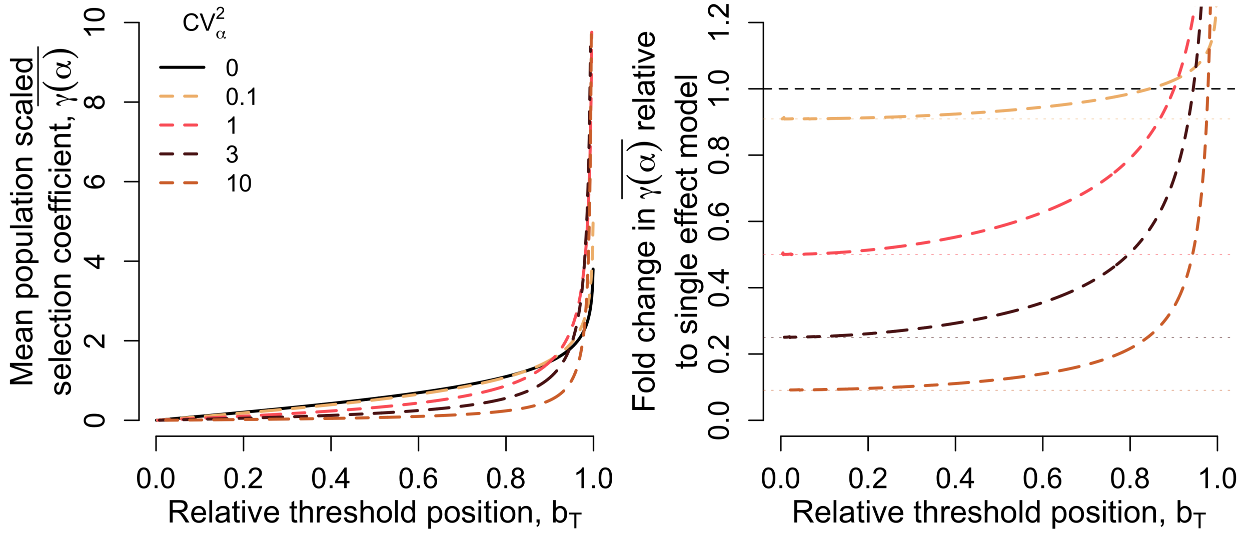

Figure S14: A) The mean scaled selection coefficient as a function of $b_T$, assuming that the effect sizes follow a gamma distribution. B) The same as in panel (A), but plotted relative to the single effect case (i.e. $CV_a^2 = 0$.)

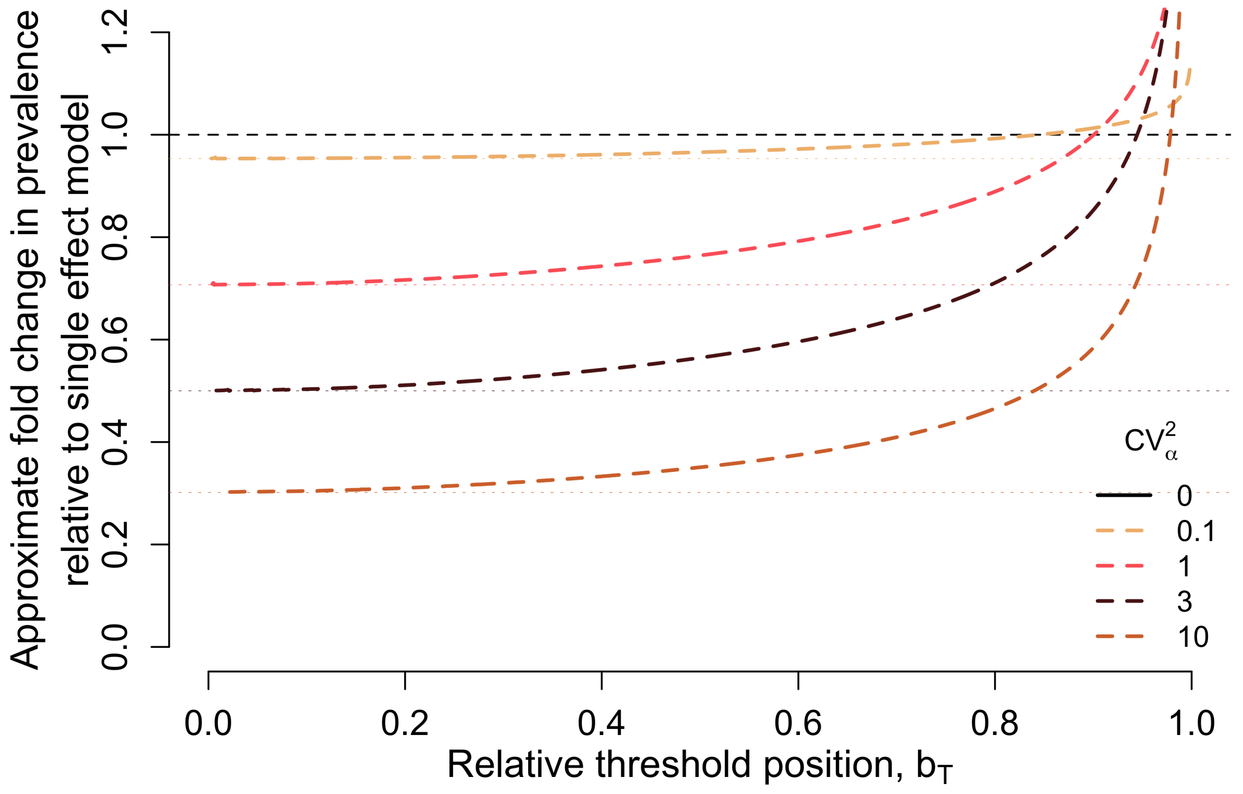

Figure S15: The ratio of the standardized threshold density in the model with a distribution of effects relative to that in the corresponding single effect model

### S7 Two Effect Model

Here, we detail our method for solving the two effect model.

In this model, we assume that a fraction $p_S = 1 - p_L$ of sites have small effects, with effect size $a_S = 1$, while the remaining fraction $p_L$ of sites have large effects with effect size $a_L \gg 1$, such that $\gamma(a_L) \gg 1$.

We model the distribution of liability in the population as a convolution of two distributions. The first is a Normal distribution, which has variance $V_{A,S} + V_E$, where $V_{A,S}$ is the contribution of small effect sites to genetic variance, and $V_E$ is the environmental variance.

The second component of our approximation for the two effect model is the distribution on the number of large effect alleles an individual carries. At each site, the mean frequency is $u/(\delta_R(a_L)C) \ll 1$, so we can model the number of large effect alleles per individual as Poisson with mean

$$\lambda_L = \frac{2Lp_L u}{\delta_R(a_L) C}. \tag{S127}$$

Given these two components, the probability that an individual's total liability exceeds the mean liability by $Z$ or more is

$$F(Z) = \sum_i P(i \mid \lambda_L) \, Q(Z \mid a_L(i - \lambda_L), V_{A,S} + V_E) \tag{S128}$$

where $Q(Z \mid \mu, \sigma^2) = 1 - \Phi(Z \mid \mu, \sigma^2)$ is the complementary CDF a Normal distribution, and $P(i \mid \lambda_L)$ is the probability that a Poisson random variable with mean $\lambda_L$ is equal to $i$. The density on total liability, in turn, is

$$f(Z) = \sum_i P(i \mid \lambda_L) \, \phi(Z \mid a_L(i - \lambda_L), V_{A,S} + V_E) \tag{S129}$$

where $\phi(Z \mid \mu, \sigma^2)$ is the PDF of a Normal distribution. (Note that equations (S128) and (S129) involve a slight abuse of notation relative to the main text, where $F(Z)$ and $f(Z)$ are defined not in terms of deviations from the mean, but in terms of absolute liability.)

We can solve this two effect model in the following way. First, because all large effect sites are fixed for the liability decreasing allele, the mutational asymmetry among the small effect sites must be

$$b(a_S) = \frac{b_T \bar{a} - p_L a_L}{p_S a_S}, \tag{S130}$$

where

$$\bar{a} = p_S a_S + p_L a_L \tag{S131}$$

is the mean effect size across all sites. Next, because the threshold density is determined entirely by the fixation dynamics of small effect sites, it is given by

$$f(T) = \frac{1}{4NCa_S} \ln \frac{1 + b(a_S)}{1 - b(a_S)}. \tag{S132}$$

This in turn allows us to compute the contribution to genetic variance from small effect sites, which is

$$V_{A,S} = \frac{2Lp_S u a_S b(a_S)}{Cf(T)}. \tag{S133}$$

Notably, to obtain equation (S133) we did not have to know anything about the large effect sites other than how many there are, how big their effects are, and the fact that they are all fixed in the protective state.

Now, the risk effect of the large effect sites is given by

$$\delta_R(a_L) = F(T^* - a_L) - F(T^*) \tag{S134}$$

where $T^* = T - \overline{Z}$ is the distance between the mean and the threshold.

Equation (S134) can be used, in conjunction with the other equations in this section, to solve for the equilibrium: given a proposed value for $\delta_R(a_L)$, we can plug $T^*$ in for $Z$ in equation (S129), set it equal to (S132)

$$\frac{1}{4NCa_S} \ln \frac{1 + b(a_S)}{1 - b(a_S)} = \sum_i P(i \mid \lambda_L) \, \phi(T^* \mid a_L(i - \lambda_L), V_{A,S} + V_E) \tag{S135}$$

and solve numerically for $T^*$. Once we have $T^*$, the right hand side of equation (S134) is fully specified. Thus, we can solve the model by continuing to update our proposal for $\delta_R(a_L)$ via a line search, until we find the value that satisfies equation (S134).

The above algorithm imagines that we specify a fixed value of $V_E$ in our solution. We found it useful to solve the model conditional on a particular heritability. To do this, we simply add one additional step. For each proposed $\delta_R(a_L)$, we compute the large effect contribution to variance in liability using our Poisson assumption as

$$V_{A,L} = a_L^2 \lambda_L \tag{S136}$$

and then compute the environmental variance as

$$V_E = \frac{1 - h^2}{h^2} (V_{A,S} + V_{A,L}) \tag{S137}$$

and solve equation (S135) as outlined above.

### S8 Simulation details

We simulated the equilibrium behavior of our model using SLiM version 3.6. Here we describe details of our simulation scheme that are not described in the main text or figure captions. The full SLiM scripts are included in the supplementary materials.

#### S8.1 Single effect simulations

A key feature of our model is that an individual's fitness depends on their absolute liability, and that mutation is state-dependent. To enforce this state dependence, we used SLiM's nucleotide model. In the single effect simulations, we encode low liability alleles as A nucleotides and high liability alleles as G nucleotides, and set the liability effect of a G allele to $a = 1$. Thus, in the single effect simulations, an individual's genetic liability is equal to the number of G alleles that they carry (including both fixed and segregating sites). We found that this nucleotide scheme in SLiM was mostly sufficient to enforce the finite-sites state-dependent mutation model. However, because of the way that SLiM internally tracks the list of sites that are segregating vs fixed, we found that occasional back mutations at derived segregating sites (e.g. a mutation from G to A at a segregating derived G allele) could create problems with our liability accounting due to a lack of clarity over whether a given site was still segregating the original mutation or not. To remedy this issue, we wrote custom Eidos code to remove such back mutations from SLiM's mutation list whenever they occur (while keeping the underlying nucleotide change).

For each simulation with a given combination of $L$ and $b_T$, we set the threshold value to $T = L(1 - b_T)$ and initialized the simulation with a population of individuals who were fixed for the liability increasing G allele at $T - 4N\Delta_U\overline{G} = L(1 - b_T(1 + 2\theta))$ sites. We then ran a burn-in of $10N$ generations before we began recording the state of the population. These choices ensure that the population will have had the opportunity to evolve neutrally and accumulate genetic variation for approximately $4N$ generations before it comes into contact with the threshold. We found that this was necessary because if we initialized the population too close to the threshold, then very little segregating genetic variance would have accumulated by the time the population came into contact with the it, so that the initial selection response was insufficient to halt the increase in liability due to the mutational pressure. This resulted in entire population pushing past the threshold shortly after coming into contact with it, at which point all individuals would have the disease and selection could no longer be effective.

By starting the population $4N$ generations-worth of mutational increments away from the threshold, we ensured that the population could accumulate levels of genetic variance close to those expected under neutrality. This is more than the amount of variance expected at MSDB, and thus sufficient to halt the increase in liability due to mutation. In practice, this was only an issue when $L = 1.5 \times 10^8$ and $b_T$ was relatively large, but we implemented it for all single effect simulations for simplicity. It is likely that even in the cases where $L$ and $b_T$ were large, starting a full $4N$ generations-worth of mutational increments away from the threshold was probably more than was necessary, but we did not find it necessary to further optimize the initialization.

After these first $4N$ generations of burn-in, the population then evolves at MSDB for approximately $6N$ generations after coming into contact with the threshold. This is long enough that essentially all segregating variation in the population once we begin recording should reflect the MSDB equilibrium.

Beyond these special considerations, we set the recombination rate among adjacent base pairs equal to $100\times$ the mutation rate to ensure that individual segregating sites are effectively independent in their evolution.

#### S8.2 Two effect simulations

The set up for our two effect simulations is nearly identical, except the obvious addition of a second class of effect sizes. To model the "large effect" sites, we further leveraged SliM's nucleotide model, using C to encode the low liability allele and T to encode the high liability allele at each of the large effect sites. To initialize each two-effect simulation, we set all large effect sites to be fixed for the low liability C allele, consistent with MSDB expectation for strongly selected sites.

We initialized the small effect sites to be fixed for the liability increase G allele at a total of $T - 4\sqrt{V_E}$ sites, so that the population would come into contact with the threshold relatively quickly, thus preventing any large effect alleles from fixing before selection could become effective. Notably, this offset of $4\sqrt{V_E}$ is much smaller than the $4N\Delta_U\overline{G}$ offset we used for the single effect simulations but did not cause problems because we did not run two-effect simulations with combinations of $L$ and $b_T$ that were both large.

#### S9 Smile plots

To generate the smile plots in Figure 6, we downloaded GWAS summary statistics for four different disease traits. We downloaded summary statistics for type 2 diabetes [12] and inflammatory bowel disease [6], and cardiovascular disease [5] from the GWAS Catalog [4] (accessions GCST010118, GCST003043, and GCST90038595, respectively), while the schizophrenia data [13] were downloaded from the Psychiatric Genomics Consortium website (https://pgc.unc.edu/for-researchers/download-results/). We then filtered all datasets to include only variants with minor allele frequency (MAF) $\geq 0.01$ and p-values $< 5 \times 10^{-8}$.

To identify independent variants, we used predefined and publicly available approximately independent linkage disequilibrium (LD) blocks. For each GWAS, we used LD blocks that we generated in a sample that was genetically similar to the GWAS sample. The type 2 diabetes GWAS that we used [12] was performed in individuals of east Asian ancestry, so for this GWAS we used the 'EAS' LD blocks generated by [7] using patterns of LD in the Han Chinese in Beijing, China (CHB) 1000 genomes sample [1, 15]. For the other three disease phenotypes, the GWAS was performed in individuals of European ancestry, so we used the 'EUR' LD blocks generated by [2]. Within each LD block, we selected the variant with the smallest p value as a representative for the block (subject to the constrait that it pass the (MAF) $\geq 0.01$ and p-values $< 5 \times 10^{-8}$ filter mentioned above). For each disease, we then plot the effect of the risk increasing allele against its frequency for each of the representative variants remaining after this filtering step.
